# A High-Resolution In Vivo Atlas of the Human Brain’s Benzodiazepine Binding Site of GABA_A_ Receptors

**DOI:** 10.1101/2020.04.10.035352

**Authors:** Martin Nørgaard, Vincent Beliveau, Melanie Ganz, Claus Svarer, Lars H Pinborg, Sune H Keller, Peter S Jensen, Douglas N. Greve, Gitte M. Knudsen

**Affiliations:** Neurobiology Research Unit & CIMBI, Copenhagen University Hospital, Rigshospitalet, Copenhagen, Denmark; Institute of Clinical Medicine, University of Copenhagen, Copenhagen, Denmark; University of Copenhagen, Department of Computer Science, Copenhagen, Denmark; Athinoula A. Martinos Center for Biomedical Imaging, Massachusetts General Hospital, Harvard Medical School, Boston, MA, USA; Medical University of Innsbruck, Department of Neurology, Innsbruck, Austria; Department of Clinical Physiology, Nuclear Medicine and PET, Righospitalet, Copenhagen, Denmark

**Keywords:** GABA, PET, Atlas, Autoradiography, mRNA, benzodiazepine binding site

## Abstract

Gamma-aminobutyric acid (GABA) is the main inhibitory neurotransmitter in the human brain and plays a key role in several brain functions and neuropsychiatric disorders such as anxiety, epilepsy, and depression. The binding of benzodiazepines to the benzodiazepine receptor sites (BZR) located on GABA_A_ receptors (GABA_A_Rs) potentiates the inhibitory effect of GABA leading to the anxiolytic, anticonvulsant and sedative effects used for treatment of those disorders. However, the function of GABA_A_Rs and the expression of BZR protein is determined by the GABA_A_R subunit stoichiometry (19 genes coding for individual subunits), and it remains to be established how the pentamer composition varies between brain regions and individuals.

Here, we present a quantitative high-resolution in vivo atlas of the human brain BZRs, generated on the basis of [^11^C]flumazenil Positron Emission Tomography (PET) data. Next, based on autoradiography data, we transform the PET-generated atlas from binding values into BZR protein density. Finally, we examine the brain regional association with mRNA expression for the 19 subunits in the GABA_A_R, including an estimation of the minimally required expression of mRNA levels for each subunit to translate into BZR protein.

This represents the first publicly available quantitative high-resolution in vivo atlas of the spatial distribution of BZR densities in the healthy human brain. The atlas provides a unique neuroscientific tool as well as novel insights into the association between mRNA expression for individual subunits in the GABA_A_R and the BZR density at each location in the brain.

## INTRODUCTION

Gamma-aminobutyric acid (GABA) is the main inhibitory neurotransmitter in the human brain and is a key component in several brain functions and in neuropsychiatric disorders, including anxiety, epilepsy, and depression [1]. The GABA_A_ receptor (GABA_A_R) is a pentameric complex that functions as a ligand-gated chloride ion channel and it is the target of several pharmacological compounds such as anesthetics, antiepileptics, and hypnotics.

Benzodiazepines are well-known for their sedative, anticonvulsive and anxiolytic effects. They act as positive allosteric modulators via the benzodiazepine binding sites (BZR) which are located between the α_1,2,3,5_ and γ_1-3_ subunits in the pentameric constellation of the postsynaptic ionotropic GABA_A_R. In total, 19 subunits (α_1-6_, β_1-3_, γ_1-3_, ρ_1-3_, δ, ε, θ, π) have currently been identified, and the constellation of subunits to form GABA_A_Rs, and consequently the expression of and affinity to the BZR site, has been shown to differ between individuals, between brain regions, and across the life span [1].

High-resolution quantitative human brain atlases of e.g., receptors represent a highly valuable reference tool in neuroimaging research to inform about the in vivo 3-dimensional distribution and density of specific targets. On the basis of high-resolution PET neuroimaging data from 210 healthy individuals, Beliveau et al. 2017 [2] created a quantitative high-resolution atlas of the serotonin system (four receptors, and the serotonin transporter), providing a valuable reference for researchers studying human brain disorders, or effects of pharmacological interventions on the serotonin system that may subsequently down-or upregulate the receptors. Because the functional organization of receptor systems may be different than commonly defined structural organizations (e.g. Brodmann [3]), it is important to define and use the organization of the receptor systems modulating brain function not only to capture a more refined view of the brain [4], but also to provide insights in novel brain parcellations.

Here, we present the first quantitative PET-based high-resolution in vivo 3D human brain atlas of the distribution of BZRs; an atlas to serve as a reference in future studies. Based on autoradiography data, we transform the PET generated atlas into brain regional protein densities of BZR and compare it to the mRNA expression for individual subunits in the GABA_A_R, using the Allen Human Brain Atlas [5].

## METHODS

### Study participants and neuroimaging

Sixteen healthy participants (9 females; mean age ± SD: 26.6 ± 8 years, range 19-46 years) were included in the study [6]. The study was approved by the Regional Ethics Committee (KF 01280377), and all subjects provided written informed consent prior to participation, in accordance with The Declaration of Helsinki II. Data from 10 individuals have entered a previous paper (Feng et al. 2016 [7]) where more detailed accounts of the methods can be found. The participants were scanned between 1 and 3 times with the High-Resolution Research Tomograph (HRRT, CTI/Siemens) PET scanner at Rigshospitalet (Copenhagen, Denmark) with the radioligand [^11^C]flumazenil (26 unique PET scans in total). After an initial transmission scan [8], the radioligand was given either as an intravenous bolus injection or as a bolus-infusion, and a PET emission scan was conducted over 90 minutes, starting at the time of injection. PET data was reconstructed into 35 frames (6×5, 10×15, 4×30, 5×120, 5×300, 5×600 seconds) with isotropic voxels of 1.2 mm using a 3D-ordered subset expectation maximum and point spread function modelling (3D-OSEM-PSF) (16 subsets, 10 iterations) with [^137^Cs]transmission scan-based attenuation correction and no postfiltering [9–11]. Arterial sampling was also performed and the arterial input function was corrected for radiometabolites [7]. The reconstructed PET data were motion corrected using the reconcile procedure in AIR (v. 5.2.5, http://loni.usc.edu/Software/AIR), and our criterion for acceptable motion was a median movement less than 3 mm across frames. The PET tracer injection protocols carried out for each subject are listed in Table S1 in the supplementary.

An isotropic T1-weigthed MP-RAGE was acquired for all participants (matrix size = 256 × 256 × 192; voxel size = 1 mm; TR/TE/TI = 1550/3.04/800 ms; flip angle = 9°) using either a Magnetom Trio 3T or a 3T Verio MR scanner (both Siemens Inc.). Furthermore, an isotropic T2-weighted sequence (matrix size 256 × 256 × 176; voxel size = 1 mm; TR/TE = 3200/409 ms; flip angle = 120°) was acquired for all participants. All acquired MRI’s were corrected for gradient nonlinearities [12] and examined to ensure absence of structural abnormalities.

The MR data was processed using FreeSurfer (v.6.0) [13], and co-registered to the PET data using a rigid transformation and a normalized mutual information cost function.

Regional time-activity curves (TACs) of the target concentration, C_T_, were obtained using PETsurfer [14] defined by the Desikan-Killiany atlas. Voxel-wise TACs in common volume space (MNI152) were obtained using the Combined Volumetric and Surface (CVS) registration, and volume smoothed with a 5 mm FWHM Gaussian filter. Surface-based TACs in common space (fsaverage) were obtained using the cortical-surface registration provided by FreeSurfer and surface smoothed by a Gaussian filter with 10 mm FWHM [13, 14].

### Quantification of BZR Availability

The PET data were quantified to estimate total distribution volumes (V_T_) for each region using steady-state analysis for the bolus-infusion experiments and Logan analysis [15] for the bolus injections, both using the metabolite-corrected plasma curve as input function (Figure S1) [7].

#### Blood Acquisition and Analysis

For the bolus-infusion experiments, blood sampling was carried out by cannula insertion into one of the cubital veins, while another cannula was inserted to the other cubital vein for radiotracer administration.

For the bolus experiments, arterial blood sampling was carried out by cannula insertion into the radial artery of the nondominant arm. During the first 10 minutes of the PET scan, blood measurements were counted continuously in whole-blood using an ABSS autosampler (Allogg Technology), and three samples were drawn manually to calibrate the autosamples. The plasma to whole-blood ratio was obtained by manually drawing blood samples at 18 time points, measured using a well counter (Cobra 5003; Packard Instruments), and decay-corrected to the time of injection of the radiotracer. For correction of radiometabolites, nine blood samples were drawn during the PET scan and analysed for metabolites using radio-high performance liquid chromatography (HPLC). The metabolite-corrected plasma input function, C_P_, was estimated as the product of the parent compound fitted using a biexponential function and the total plasma concentration.

#### Data Analysis

An overview of the complete analysis workflow can be found in Figure S1 in the supplementary. The metabolite-corrected blood data curves were interpolated to the time points of the PET scan using a cubic spline. All resulting blood curves used for kinetic modeling can be found in the supplementary material.

For bolus-infusion experiments, the plateau for steady-state was used to calculate V_T_ as an average of the ratio between C_T_ and C_P_ from 35 to 80 minutes after the beginning of tracer administration [7]. For the bolus injections, the invasive Logan was used to estimate V_T_ using the metabolite-corrected input function with t*=35 minutes for all regions and subjects [16]. Since the non-displaceable distribution volume, V_ND_, is small but not negligible for [^11^C]flumazenil [16], the outcome of these analyses, V_T_, reflects the sum of specific (i.e., BZR) bound distribution volume, V_S_, and V_ND_ for all regions, including surface- and voxel maps for all scans. All kinetic models applied in this paper were implemented in MATLAB v. 2016b.

### In Vivo Binding and Autoradiography

The conversion from 3D V_T_ images with the unit ml/cm^3^ to BZR density (pmol per gram protein) was done by normalizing regional V_T_ with the corresponding regional postmortem human brain [^3^H]diazepam autoradiography data from Braestrup et al. 1977 [17]. This included 11 brain regions; inferior frontal cortex, inferior central cortex, occipital cortex, temporal cortex, cerebellum, hippocampus, amygdala, thalamus, caudate, putamen and pons. As a linear relationship between the regional average V_T_’s across subjects and the corresponding BZR densities could be established, V_ND_ could be estimated from the intercept (Figure 1B). To account for noise in the autoradiography estimates (y-axis), the standard deviation reported in [17] was used to draw random samples from a normal distribution for each region, specified by the region-specific mean and standard deviation. This procedure was repeated 1,000 times, and parameter estimates were fitted for each iteration to obtain an intercept and a slope using a linear model. A subject-specific mean estimate of the slope and intercept was obtained by averaging over the 1,000 estimates (Table S2). Finally, average V_T_ estimates were obtained by averaging across subjects, and fitted using a linear model to obtain a groupwise association between autoradiography and V_T_ (Figure 1B). This model was used to convert all surface- and voxel maps into densities of BZR availability (Figure 1A & Figure S1). Units of pmol/g tissue were converted to pmol/ml using a gray matter density of 1.045 g/ml, as done in [2].

**Figure 1:**
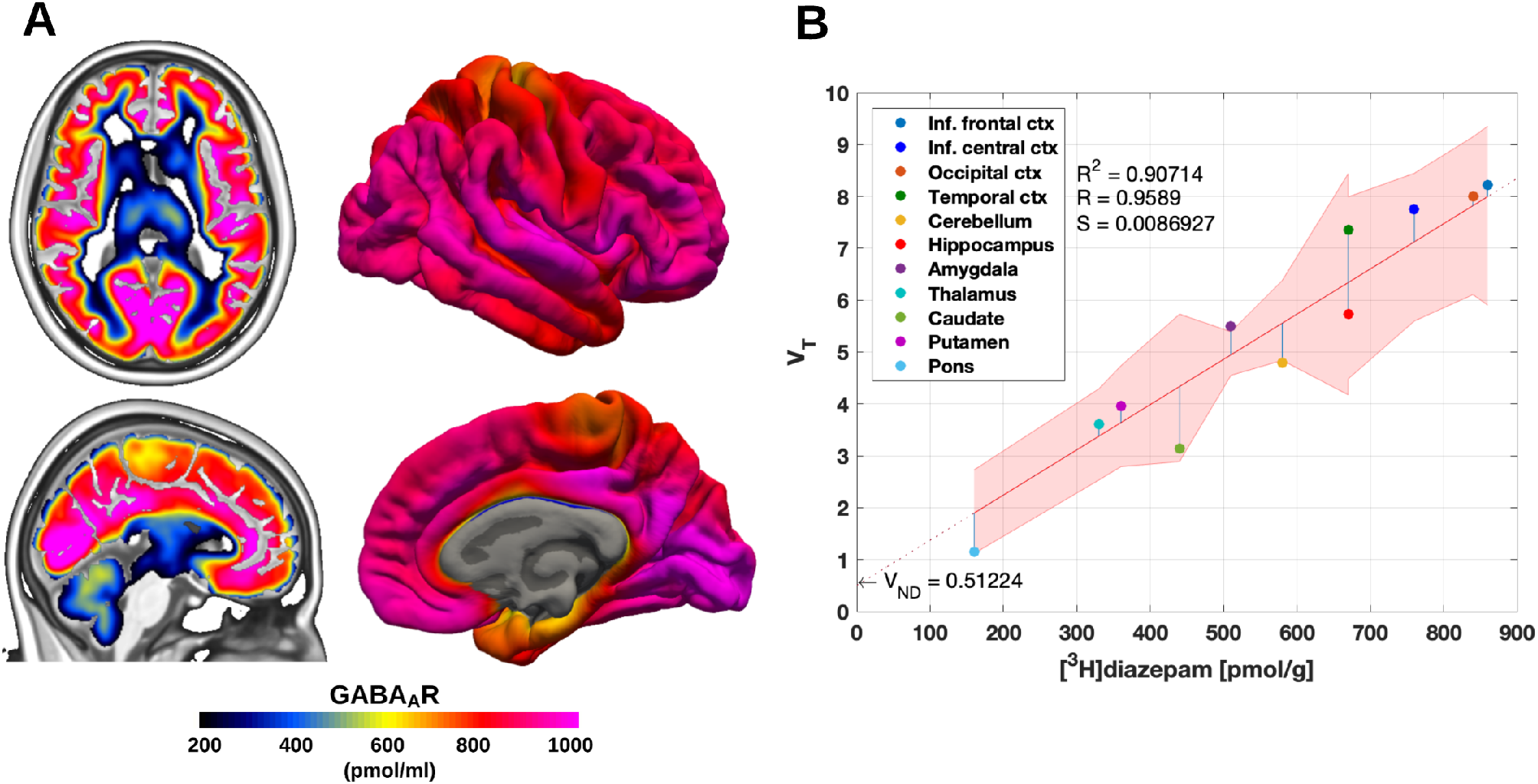
**(A)** High resolution human brain atlas of GABA_A_R density (pmol/ml) in MNI152 space (left) and in fsaverage space (right) **(B)** Average regional distribution volumes (V_T_) and benzodiazepine receptor density for the GABA_A_R. The regional V_T_’s determined by PET were matched to the corresponding regions from the [^3^H]diazepam autoradiography data. The regression is shown as the black line, and the intercept is the non-displaceable distribution volume (V_ND_). The shaded area is the 95% confidence interval.

### In Vivo Binding and mRNA Expression

Regional V_T_ estimates were compared to mRNA expression values of the 19 subunits of the GABA_A_R obtained using the Allen Human Brain Atlas (AHBA) [5]. The AHBA contains probe information from 6 human postmortem brains, where each probe provides mRNA levels (log2 intensity) for all genes and coordinates in MNI152 space. The values from each probe were matched to a brain region defined by the Desikan-Killiany atlas, and averaged within regions to provide a regional estimate of mRNA expression for each of the 19 subunits. Further information can be found in [2].

A linear model between the BZR density and the mRNA expression for each of the 19 subunits was used to examine the correspondence between regions. The intercept (I) was estimated to represent the minimally required expression of mRNA levels for each subunit to translate into BZR protein.

### Statistical Analysis

A linear mixed effects model was used to examine the effects of age, sex, administration form (bolus or bolus-infusion) and multiple scan sessions for a single subject, on the regional estimates of V_T_. The model included age, sex, and the interaction between age and sex as fixed effects. The administration form (bolus or bolus-infusion) and subjects (i.e. multiple scan sessions for a single subject) were modeled as random effects. The P-values obtained from each regional analysis (N = 11) were adjusted for multiple comparisons using false-discovery rate (FDR = 0.05).

A principal component analysis (PCA) was carried out using the mRNA expression for the 19 subunits and the BZR density for all subcortical and cortical regions (N = 48). Prior to running the PCA, the data was standardized (mean subtracted and normalized by the standard deviation). All the data is available in the supplementary material.

## RESULTS

The regional V_T_’s and the postmortem human brain autoradiography were highly correlated (Pearson’s R=0.96) and the intercept, V_ND_, was low, on average 0.51 across subjects (range 0.04-1.31) (Figure 1). Using a linear mixed effects model, we found no age or gender related effects on regional estimates of V_T_, nor an effect of administration form or multiple scan sessions for a single subject (Table S3). The BZR densities were high in cortical areas (600-1000 pmol/ml), medium in subcortical regions (300-600 pmol/ml), and low in the brainstem (<200 pmol/ml).

The association between BZR densities and mRNA expression showed a medium to high correlation for each of the subunits included in the most common pentameric GABA_A_R, i.e., α_1_β_2_γ_2_ (48% mRNA expression in the brain [1], Table I) with Pearson’s R > 0.62 (range: 0.62-0.88, P<0.0001) for all 3 subunits (Figure 2A-C).

**Figure 2:**
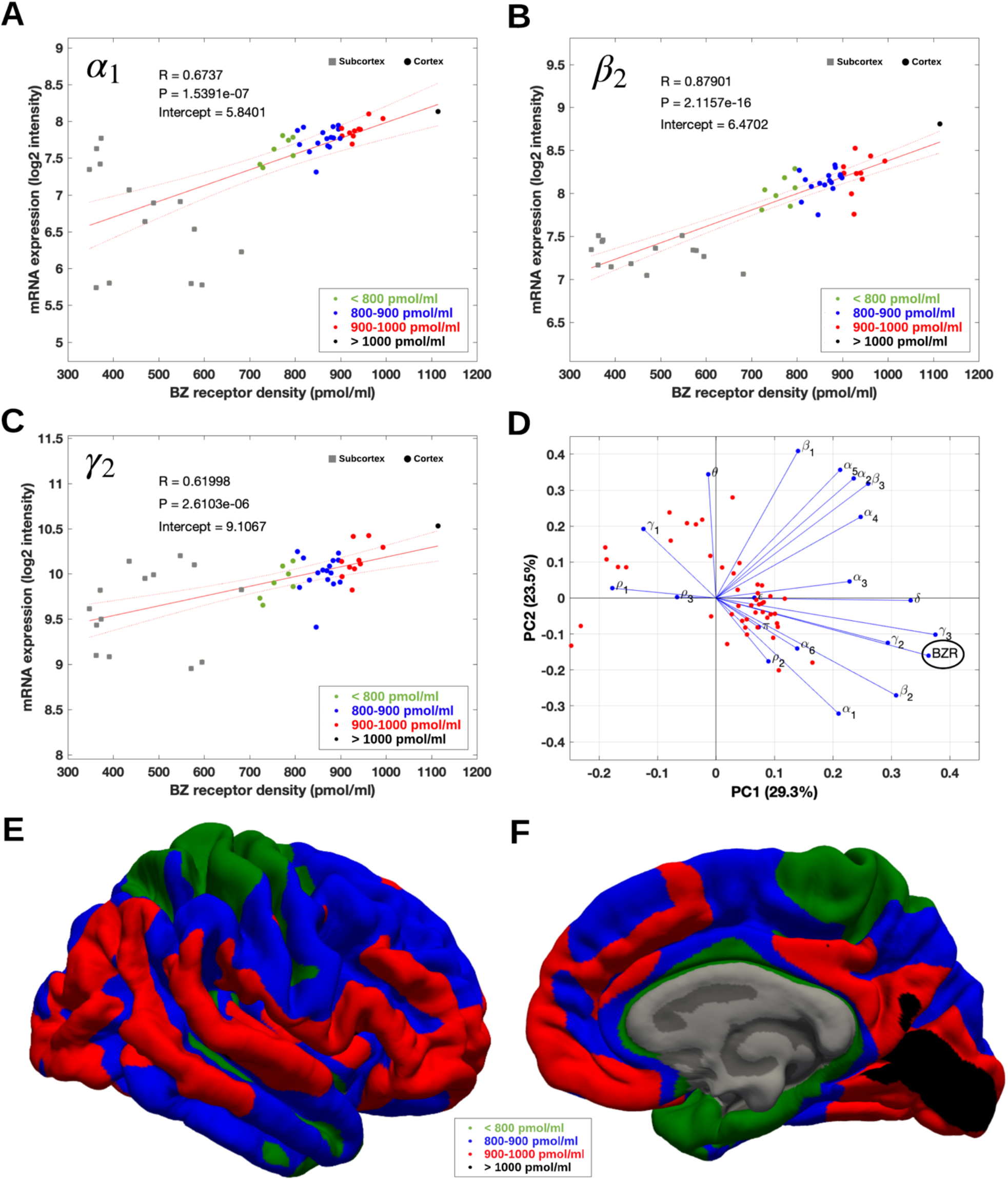
**(A-C)** Association between mRNA expression (log2 intensity) and BZR density for the subunits of the GABA_A_R most commonly represented in the BZR, α_1_ in **(A)**, β_2_ in **(B)**, and γ_2_ in **(C)**. The points are regional estimates for subcortex (squares) and cortex (round dots), and are color coded according to the density, green (<800 pmol/ml), blue (800-900 pmol/ml), red (900-1.000 pmol/ml), and black (>1.000 pmol/ml). **(D)** Biplot of the (scaled) first two principal components (% variance explained) of a PCA of the 19 subunits and BZR. **(E-F)** The spatial distribution of BZR density according to the specified color coding shown on the lateral and medial surface of the brain.

**Table I:** Association between mRNA subunit expression (log2 units) and benzodiazepine receptor density (pmol/ml) for 14 of the available 19 subunits of the GABA_A_ receptor, explaining 98.4% of the total variance across brain regions. The associations for the individual subunits are summarized by the Pearson correlation coefficient (R), the uncorrected P-value (P), the intercept (I), and the variance explained (V). The variance explained was obtained from Sequiera et al. 2019 [1]

| | $\alpha_1$ | $\alpha_2$ | $\alpha_3$ | $\alpha_4$ | $\alpha_5$ | $\alpha_6$ | $\beta_1$ | $\beta_2$ | $\beta_3$ | $\gamma_1$ | $\gamma_2$ | $\gamma_3$ | $\theta$ | $\delta$ |
| --- | --- | --- | --- | --- | --- | --- | --- | --- | --- | --- | --- | --- | --- | --- |
| R | 0.67 | 0.28 | 0.39 | 0.33 | 0.15 | 0.36 | 0.03 | 0.88 | 0.26 | -0.37 | 0.62 | 0.92 | -0.33 | 0.71 |
| P | <0.01 | 0.05 | <0.01 | 0.02 | 0.31 | 0.01 | 0.84 | <0.01 | 0.07 | 0.01 | <0.01 | <0.01 | 0.02 | <0.01 |
| I | 5.8 | 7.2 | 5.1 | 4.9 | 6.9 | 1.9 | 8.5 | 6.5 | 7.4 | 8.2 | 9.1 | 1.5 | 3.2 | 4.5 |
| V | 16 | 11 | 2 | 2 | 5 | 3 | 5 | 12 | 7 | 7 | 20 | 0.2 | 0.2 | 8 |

The PCA showed that 6 components explained >90% of the total variance, with the first and second component explaining 29.3% and 23.5%, respectively (Figure 2D). The biplot in Figure 2D shows the contribution of each subunit to the BZR density, indicating a high correlation between the BZR density and the subunits α_1_, β_2_, γ_2_, γ_3_ whereas the subunits γ_1_, ρ_1_ and ρ_3_ were negatively correlated with the BZR density.

Table I shows the linear association (Pearson’s R) between mRNA expression of 14 of the 19 available subunits of GABA_A_R and BZR density, ranging between −0.37 (γ_1_) and 0.92 (γ_3_). Six of the subunits (α_1_, α_3_, β_2_, γ_2_, δ_3_ and δ) showed a significant correlation (corrected for multiple comparisons using Bonferroni, P < 0.01). The intercept (I) for each of the subunits is the minimal mRNA expression (log2 intensity unit) required for translation into BZR protein and ranged from 1.5 (γ_3_) to 9.1 (γ_2_). Only the six subunits with a significant association (P < 0.01) between mRNA expression and BZR density (α_1_, α_3_, β_2_, γ_2_, γ_3_ and δ) were considered to have valid intercepts.

The variance explained (V) for each of the subunits was obtained from Sequiera et al. 2019 [1] and shows the proportional contribution (%) from each subunit on the total mRNA expression in the brain across all subunits. Most of the variance was explained by the subunits α_1_, α_2_, β_2_, γ_2_, and δ, contributing with 67% of the variance (Table 1).

## DISCUSSION

Here, we present the first quantitative high-resolution in vivo human brain atlas of BZR protein density, freely available at https://nru.dk/BZR-atlas. All the data and supplementary information is also publicly available.

First, we validated the association between the in vivo BZR protein density and mRNA expression. This validation showed a very strong relationship between [^11^C]flumazenil-PET and the autoradiography data from Braestrup et al. 1977 [17], supporting the use of [^11^C]flumazenil-PET as a reliable putative marker for BZR availability in the living human brain.

Autoradiography is the gold standard for estimating the density of specific receptors in the brain [17]. Braestrup et al. 1977 [17] used the agonist tracer [^3^H]diazepam to estimate the density of available BZRs, but despite that [^3^H]diazepam [18] and the antagonist tracer [^11^C]flumazenil used in PET [19, 20] have different pharmacological properties, there is no evidence for differences in their ability to bind to other receptors, i.e. they both display low non-specific binding. However, while the non-specific component is often neglected in [^11^C]flumazenil-PET studies, we observed a minor non-specific component across all subjects and scans that will potentially bias the subject-specific estimate of BZR availability if not taken into account. [^11^C]flumazenil-PET studies commonly use reference tissue modeling and the pons as reference region for estimation of BZR availability [21]. For all subjects and scans in this study, the V_T_ in pons was higher than the V_ND_, suggesting that this region has specific BZR binding. Therefore, reference tissue modeling of the PET data using pons as a reference region is not entirely valid and may be associated with an underestimation of GABA_A_R availability [21].

To make our estimates from [^11^C]flumazenil-PET more comparable to Braestrup et al. 1977 [17], we carried out a post-hoc analysis to estimate the specific binding in the occipital cortex as V_S_ = (f_p_*B_avail_)/K_D_ [15] using f_p_=0.04 [14], B_avail_ = 880 pmol/ml, K_D_ = 4.8 nM [17]. This resulted in a V_S_ equal to 7.3 ml/cm^3^. This estimate is very close to the average specific binding (Vs = 7.5 ml/cm^3^) obtained in our PET studies. In comparison, Lassen et al. 1995 [20] used [^11^C]flumazenil to estimate the B_avail_ = 120 nmol/L, K_D_ = 14 nM, and BP = B_avail_/K_D_ = 8.9 ml/cm^3^, based on partial blocking of the receptor. Several other studies have reported similar or lower B_avail_ and K_D_ estimates using PET [22, 23]. However, taking into account the free fraction in plasma of flumazenil (f_p_ = 50%), the specific binding in Lassen et al. 1977 is equal to 4.3 ml/cm^3^. The estimates by Lassen et al. 1993 [20] are much lower for both V_S_ and B_avail_ compared to the estimates obtained in this study and Braestrup et al. 1977 [17], and is likely the result of limited spatial resolution of the PET scanner (ECAT 953b; CTI PET Systems, Knoxville, TN, USA) [11], and increased partial volume effects (PVEs). Nevertheless, by using a high-resolution PET scanner as the HRRT to limit PVEs, [^11^C]flumazenil-PET with arterial blood measurements can be used as a reliable putative marker for BZR availability in the living human brain.

The second validation showed a significant association across brain regions between the expression of mRNA levels for the subunits α_1_, α_3_, β_2_, γ_2_, γ_3_ and δ, and the BZR protein density. Together, these six subunits explain 67% of the total mRNA expression across the brain. The results confirm previous evidence on the existence of a BZR binding site located between the α_1_ and γ_2_ subunits in the commonly expressed pentameric GABA_A_R subtype α_1_β_2_γ_2_ [24]. We provide novel evidence of the minimally required expression of mRNA for the translation of BZR proteins for the α_1_ (= 5.8 log2 intensity), β_2_ (= 6.5 log2 intensity) and γ_2_ (= 9.1 log2 intensity) subunits. Therefore, an imbalance in the expression of mRNA may have consequences in neuropsychiatric disorders, with the inability to produce BZR proteins due to reduced expression of mRNA. In general, the association between mRNA subunit expression and BZR density was particularly high throughout the neocortex (Figure 2), whereas the associations varied more in the subcortical regions (BZR density < 700 pmol/ml). The brain region with the highest BZR density was the occipital cortex, which has been documented to be particularly rich in both α_1_ and β_2_ subunits [1]. The remaining subunits showed a more variable association between transcription and translation, which might be caused by regional variations, due to either transport of the subunit protein away from the site of production or due to non-linear relationships between expression and translation. An indirect relationship between transcription and translation is not uncommon and has also been observed for other proteins, e.g., the serotonin transporter and the 5-HT_2A_ receptor [2].

Subcortical regions such as the striatum have a high expression of α_4_ subunits, and the amygdala has a high expression of the α_2_, β_1_ and γ_1_ subunits [1]. In our work, the association between mRNA expression of the γ_1_ subunit and BZR density was found to be negative, consistent with existing evidence that the expression of β_1_ is negatively correlated with the expression of β_2_ [1]. Together, this evidence points toward a reduced BZR density in these regions, likely caused by the stoichiometry of the subunits to form the GABA_A_R. For an in-depth analysis on the subunits only we refer the reader to Sequiera et al. 2019 [1].

Our study is not without limitations. First, the log2 intensity unit does not directly reflect quantitative mRNA expression levels, so only the relative differences in mRNA expression are useful for comparison between subunits.

Second, the association between mRNA and BZR density only allows for a direct comparison between the individual subunit and BZR density and not their interaction with other subunits. In addition, it cannot be excluded that different BZR binding pentamers have different affinity for ^11^C-flumazenil. For example, it has been shown that flumazenil has a 200-fold higher affinity to receptors with the configuration α_1_, β_2_ and γ_2_, than the affinity to receptors containing the α_6_ subunit [19]. However, as PET acquisitions are carried out using tracer doses (i.e,, ≪ K_D_) this should not impact the estimation of the specific binding of BZRs.

In the future, the development of GABA_A_R subunit-specific radioligands would be helpful to examine the GABA-ergic system in more detail. With the addition of such tools, one could gain even more insight into if pentamer subunit composition affects the affinity to pharmacological targets and if certain brain disorders are associated with abnormalities in subunit composition of the GABA_A_R.

## CONCLUSIONS

This high-resolution in vivo atlas of the spatial distribution of BZR densities in the healthy human brain provides a highly valuable tool for investigation of the GABA system, e.g., as a reference for patient groups with alterations in the cerebral GABA system. The findings provide additional insights into the association between mRNA expression for individual subunits in the GABA_A_R and the BZR density at each location in the brain and may be used to evaluate the efficacy of pharmacological targets acting on the BZR.

## ACKNOWLEDGEMENTS

We wish to thank all the participants for kindly joining the research project. We thank the John and Birthe Meyer Foundation for the donation of the cyclotron and HRRT scanner used in this study. Nic Gillings, Bente Dall, and Ling Feng are also greatfully acknowledged for their expert assistance. We also thank the VU University Medical Centre, Amsterdam, Netherlands, for sharing their knowledge and expertise on [^11^C]flumazenil.

MN was supported by the National Institutes of Health (Grant 5R21EB018964-02), the Lundbeck Foundation (Grant R90-A7722), and the Independent Research Fund Denmark (DFF-1331-00109 & DFF-4183-00627). DNG was supported by NIH grant R01EB023281 and R01NS105820.

## AUTHOR CONTRIBUTIONS

MN, VB and GMK contributed to the design of the work and analyzed the data. MN, VB, MG, CS, SHK, PSJ, LHP, DNG, and MGK interpreted the data. All authors critically reviewed the manuscript and approved the submitted version.

## SUPPLEMENTARY MATERIAL

Please find attached the supplementary material/data.

