## Supplementary figures and images for "A High-Resolution In Vivo Atlas of the Human Brain’s Benzodiazepine Binding Site of GABA_A_ Receptors"

### Cpfit.png

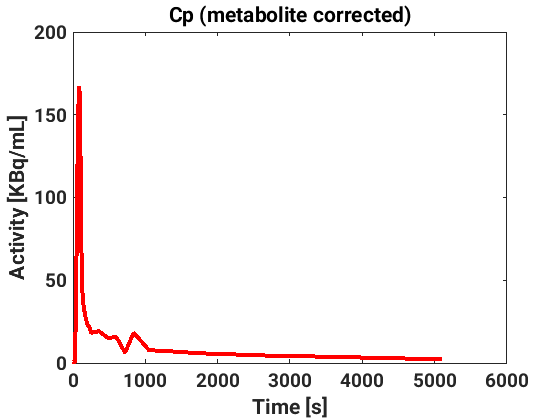

### Cpfit.png

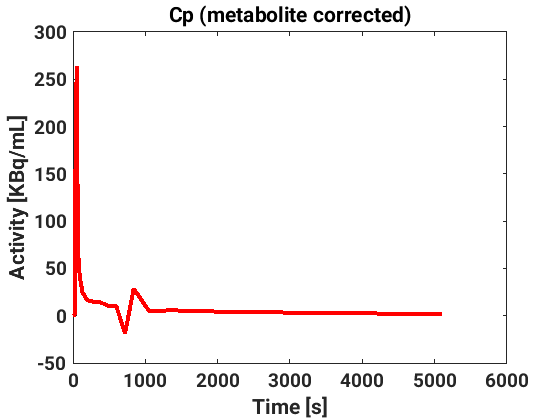

### Cpfit.png

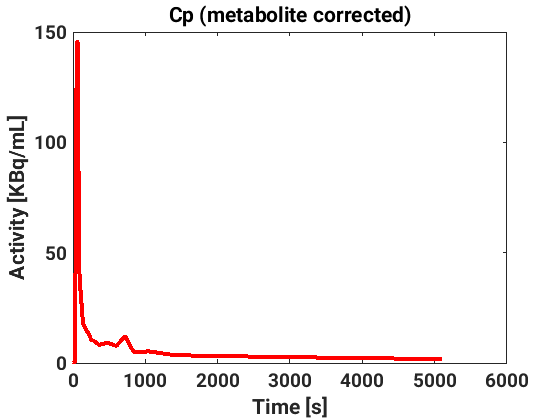

### Cpfit.png

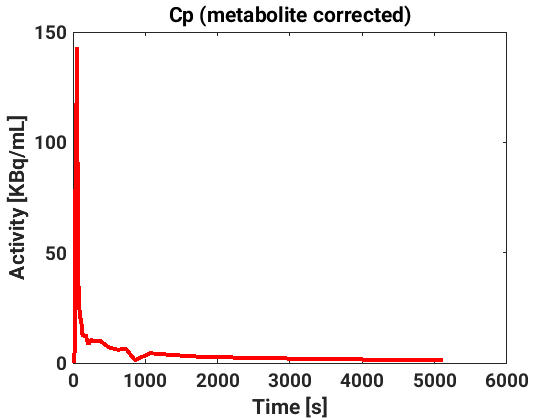

### Cpfit.png

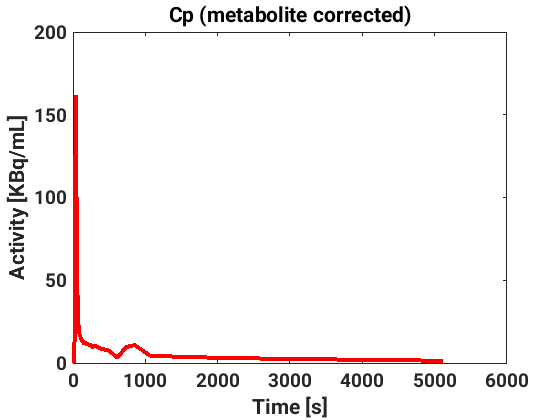

### fit_insula.png

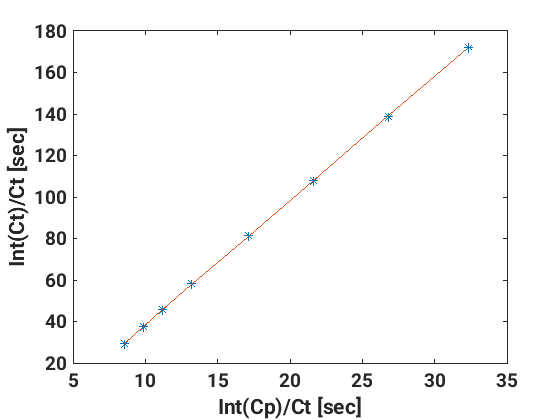

### fit_insula.png

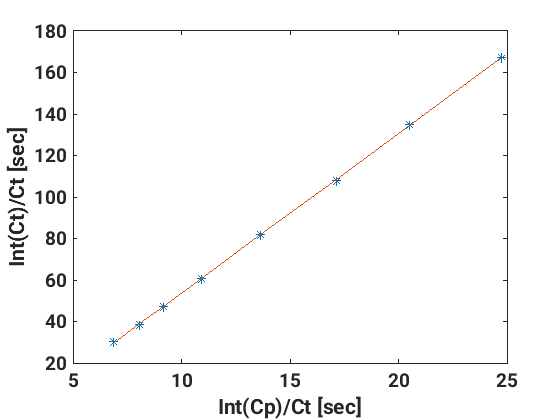

### fit_insula.png

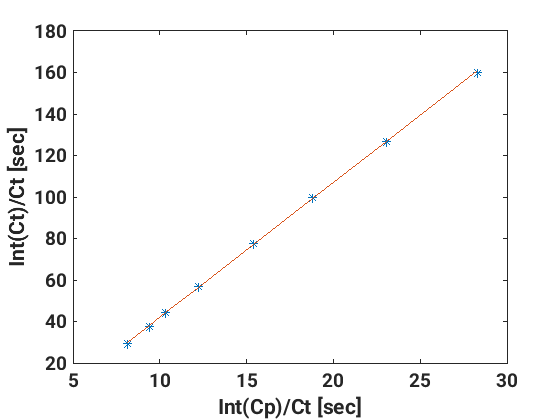

### fit_insula.png

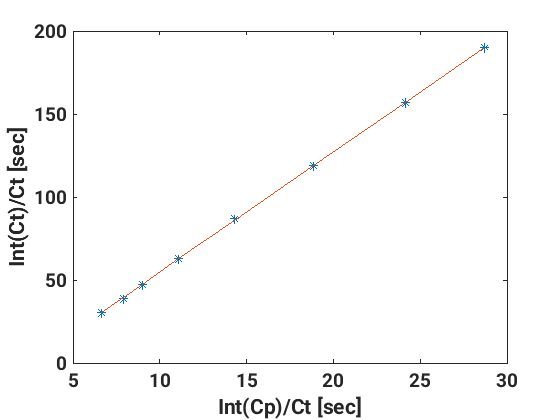

### fit_insula.png

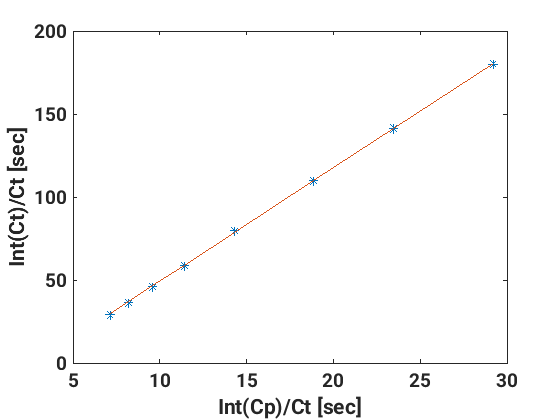

### GABRA1.png

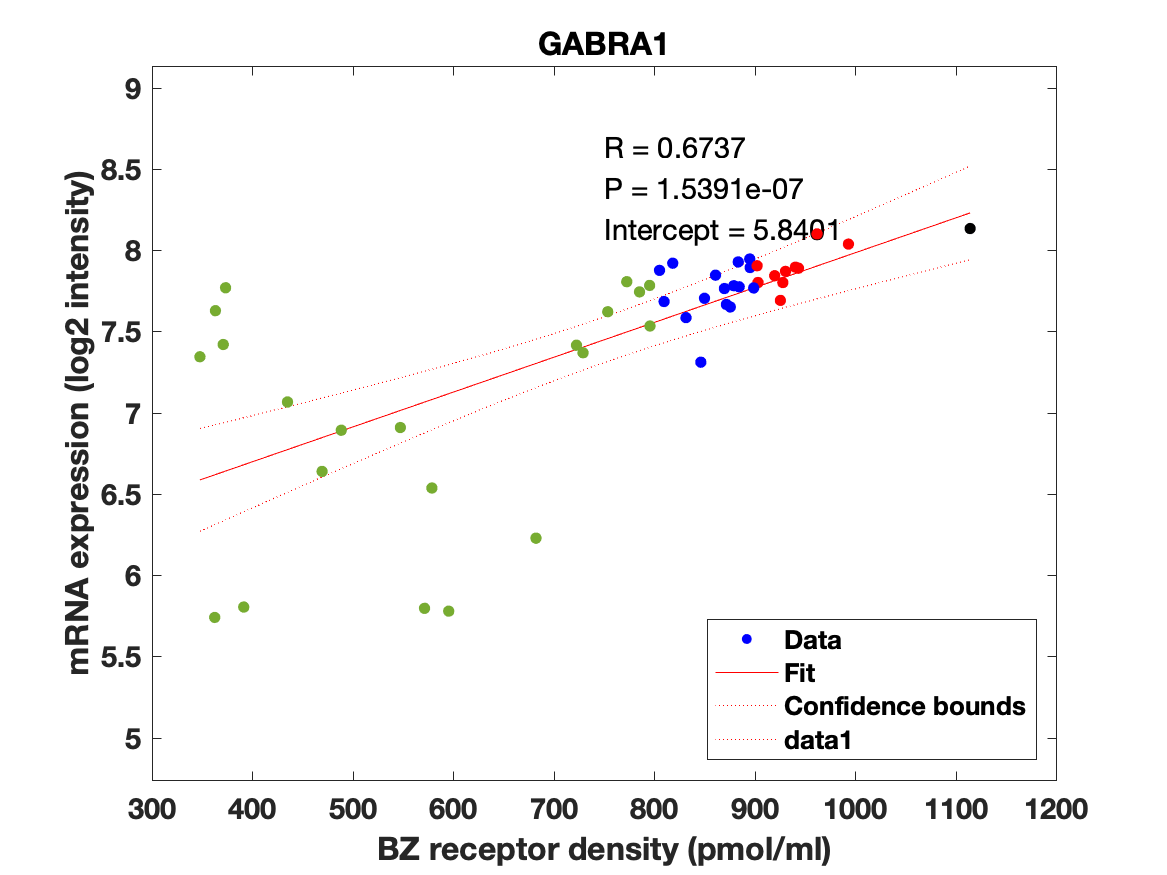

### GABRA2.png

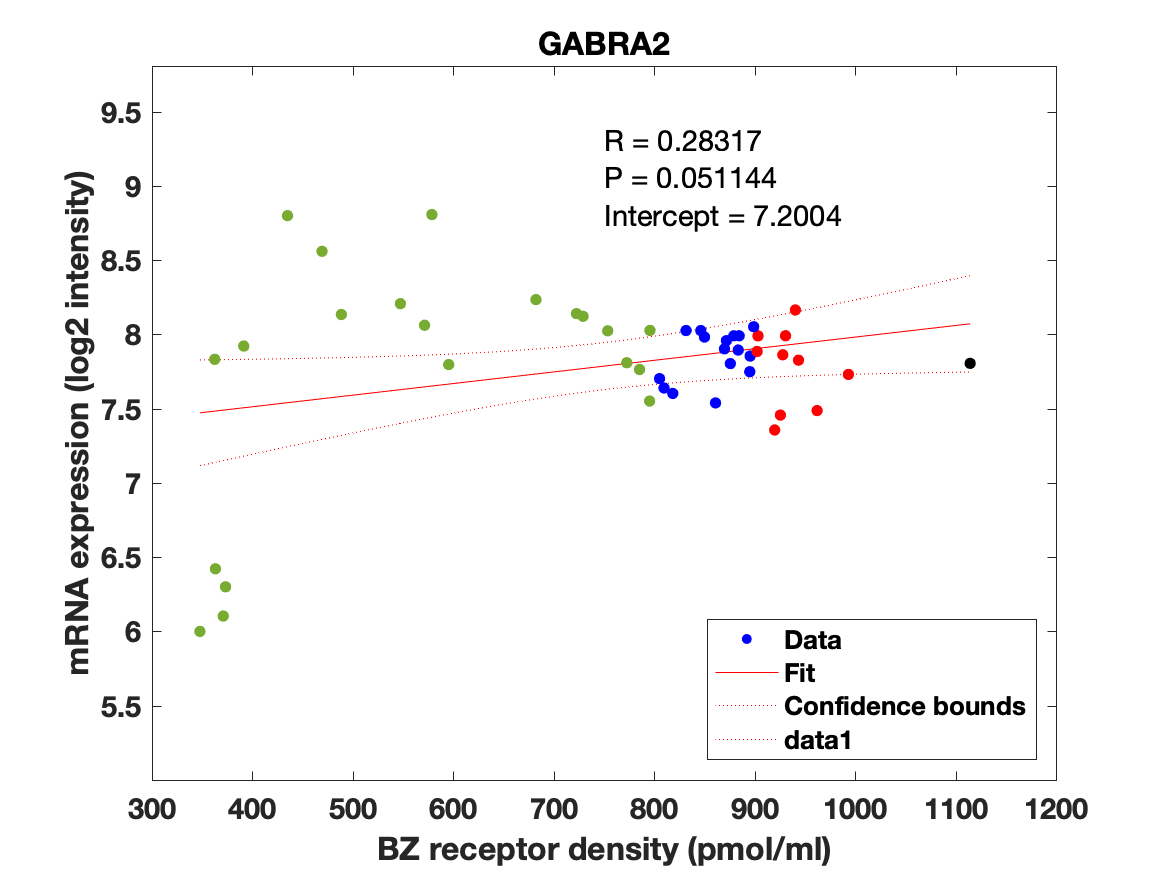

### GABRA3.png

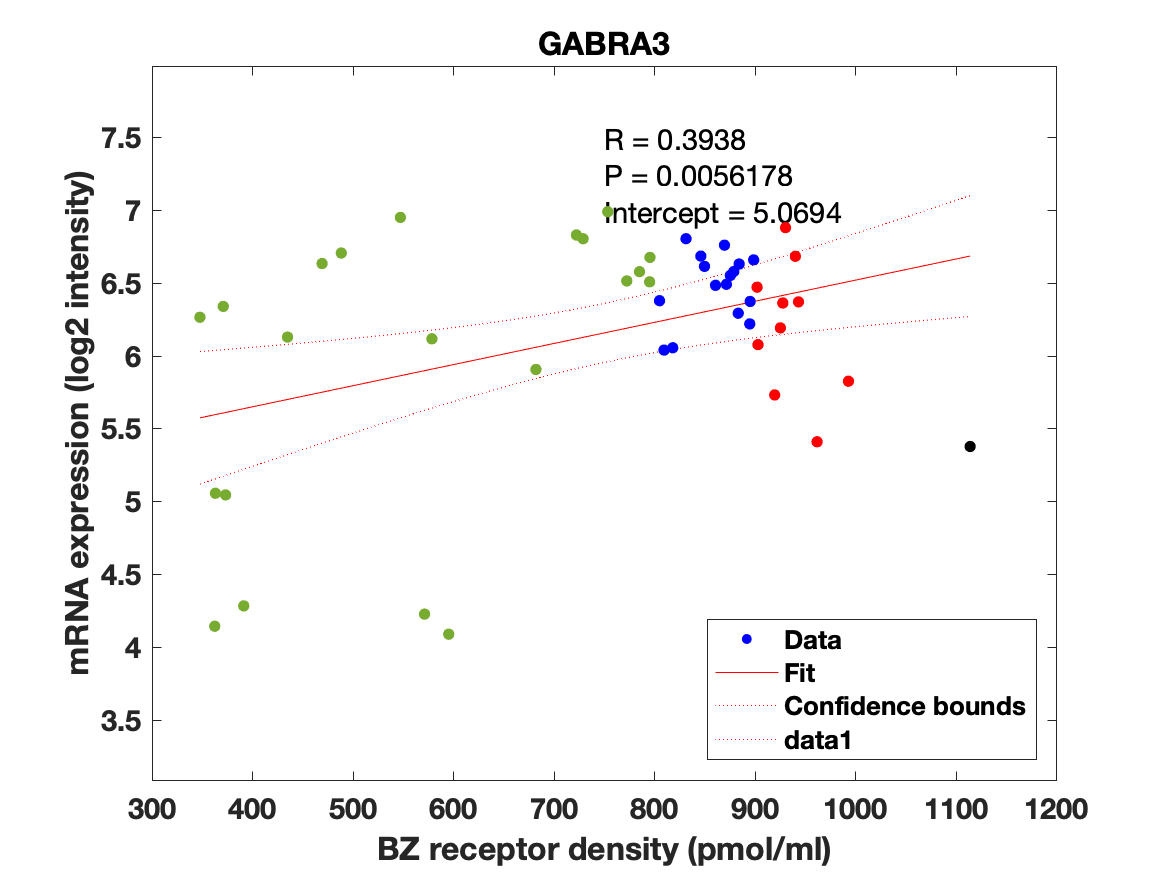

### GABRA4.png

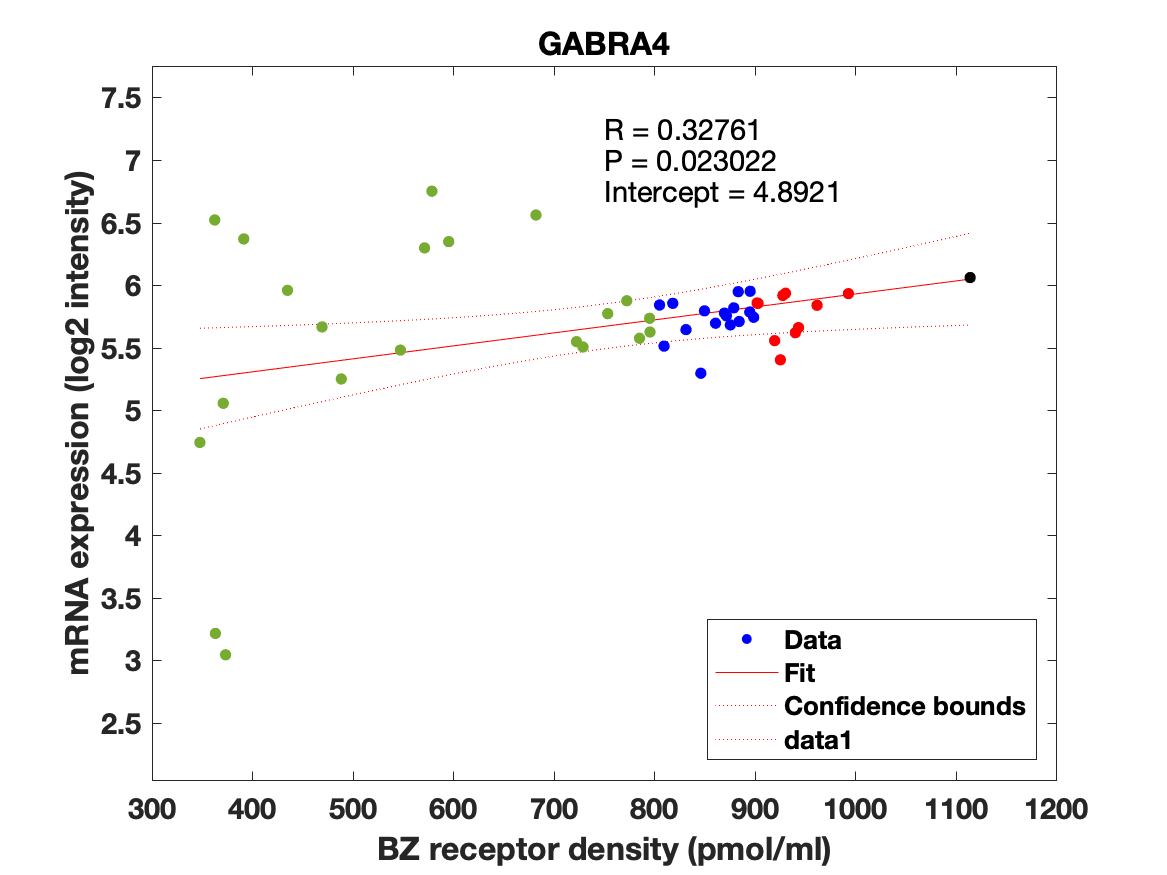

### GABRA5.png

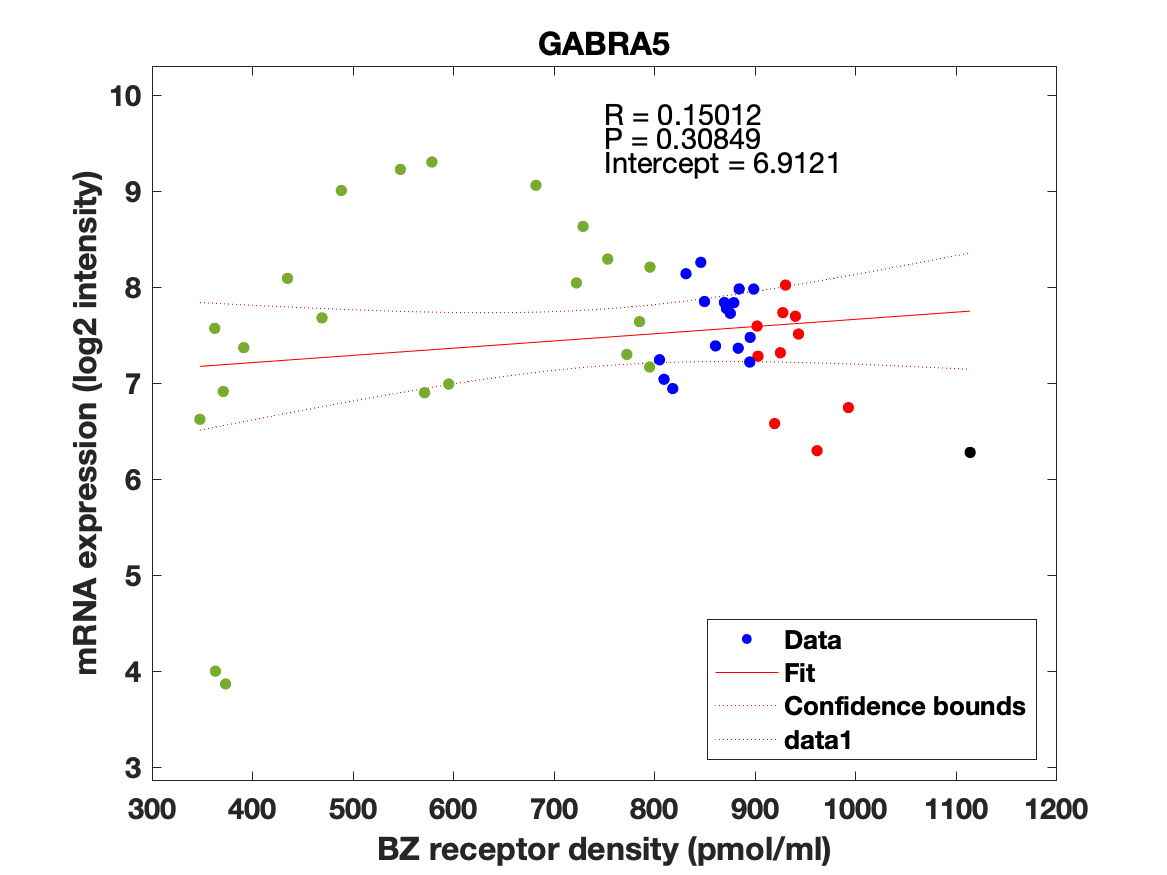

### GABRA6.png

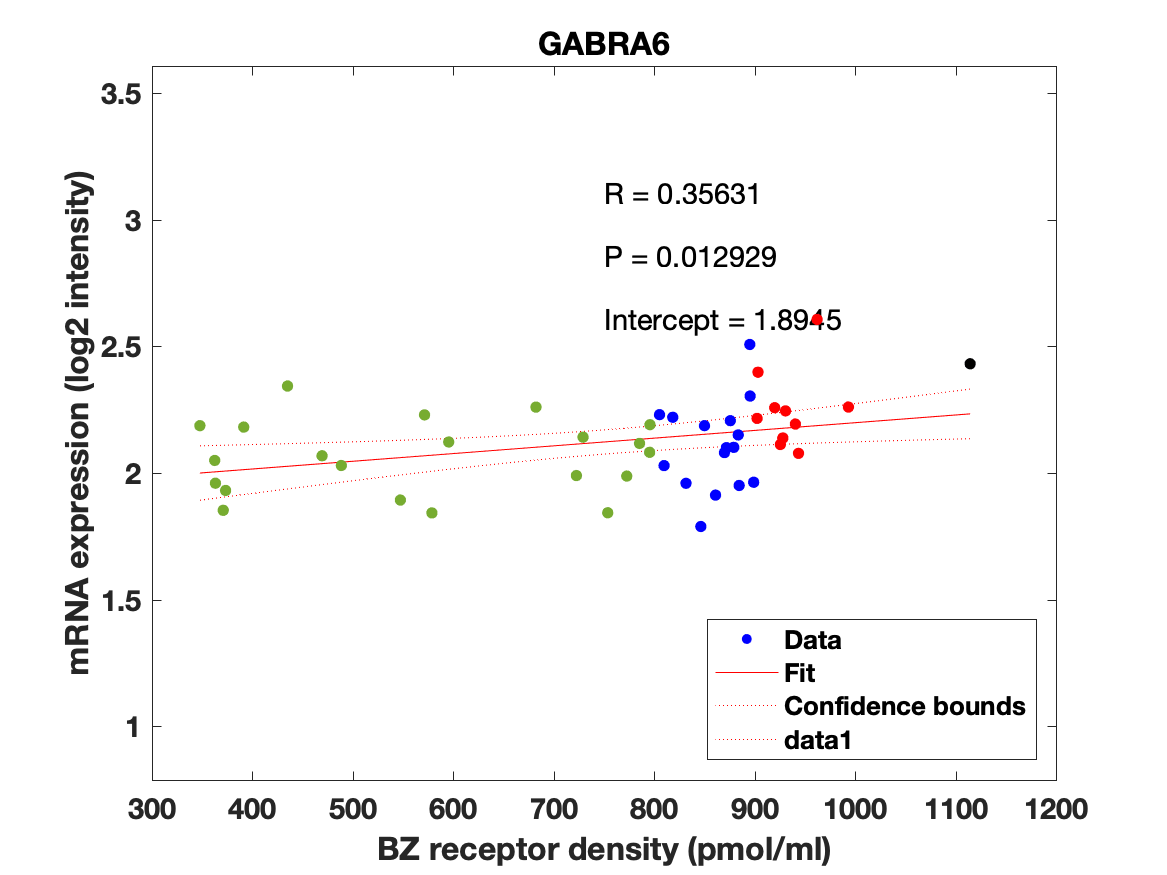

### GABRB1.png

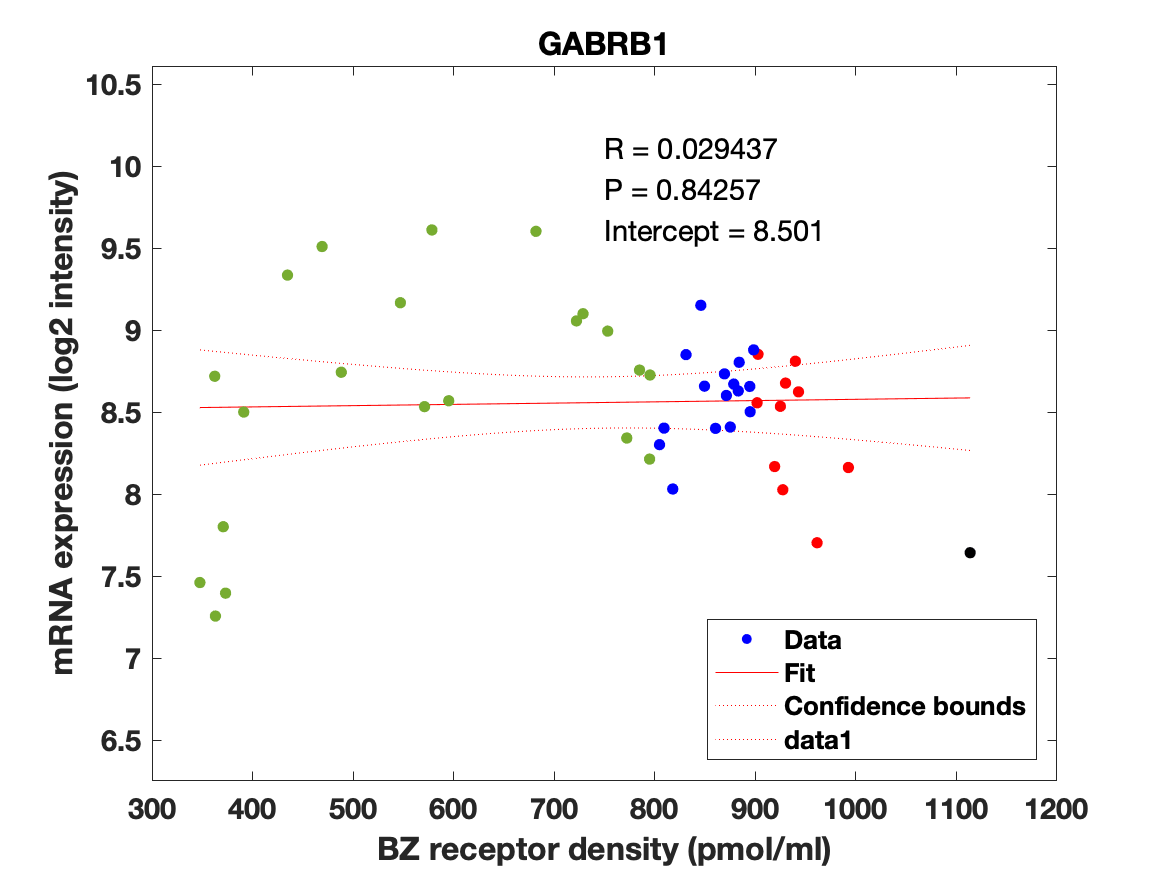

### GABRB2.png

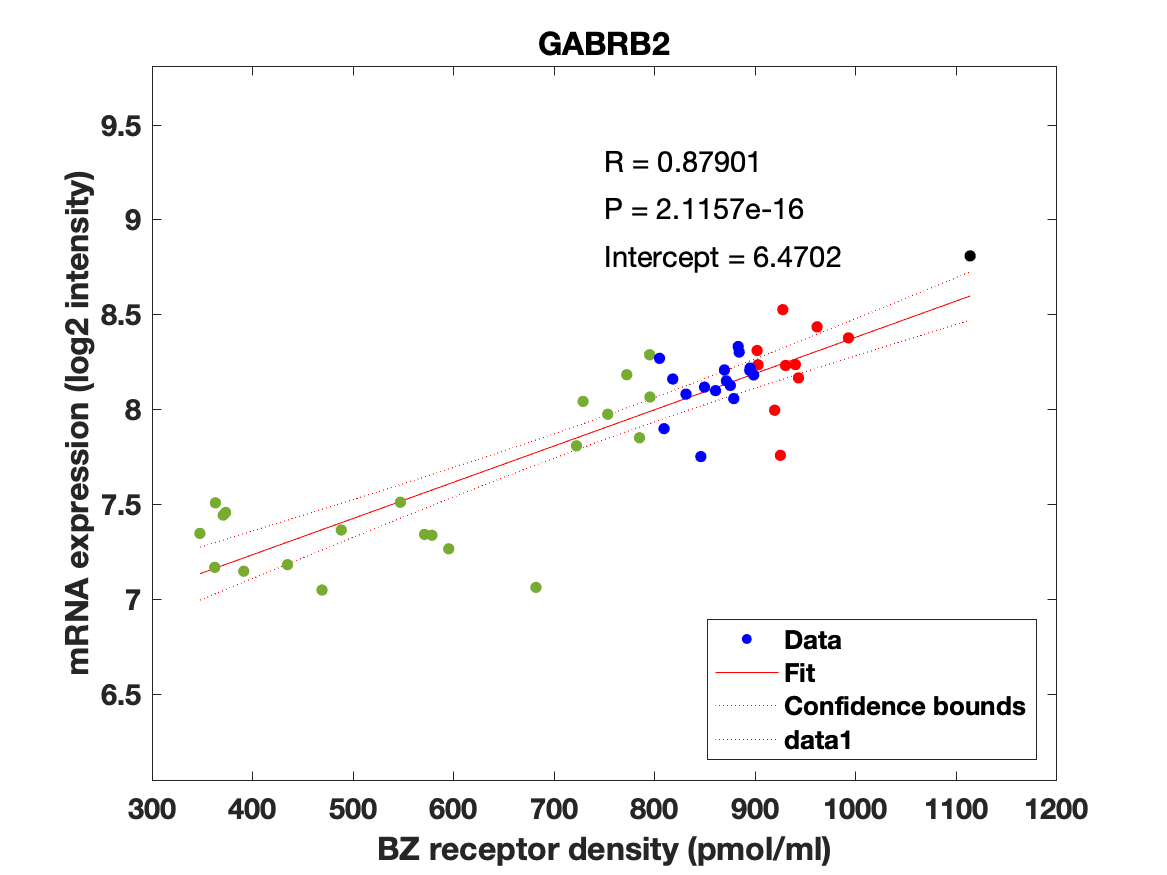

### GABRB3.png

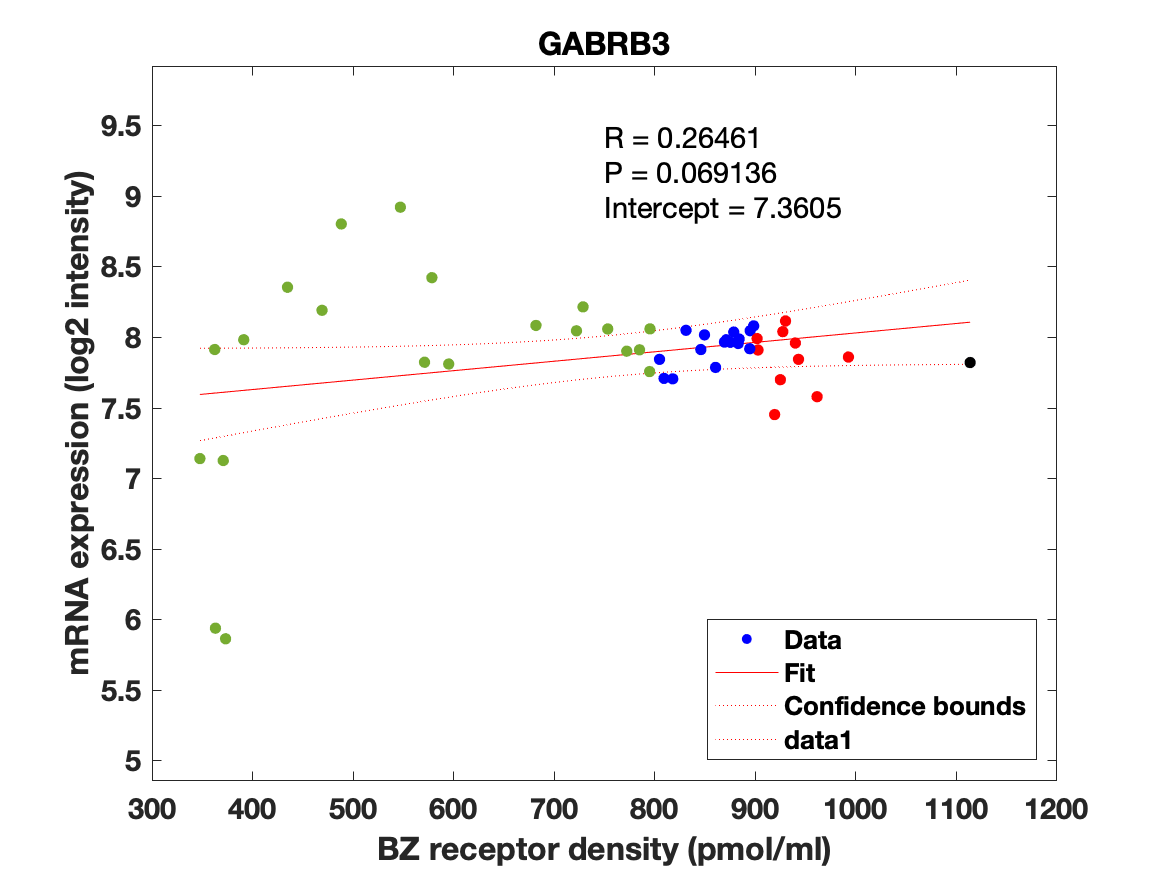

### GABRD.png

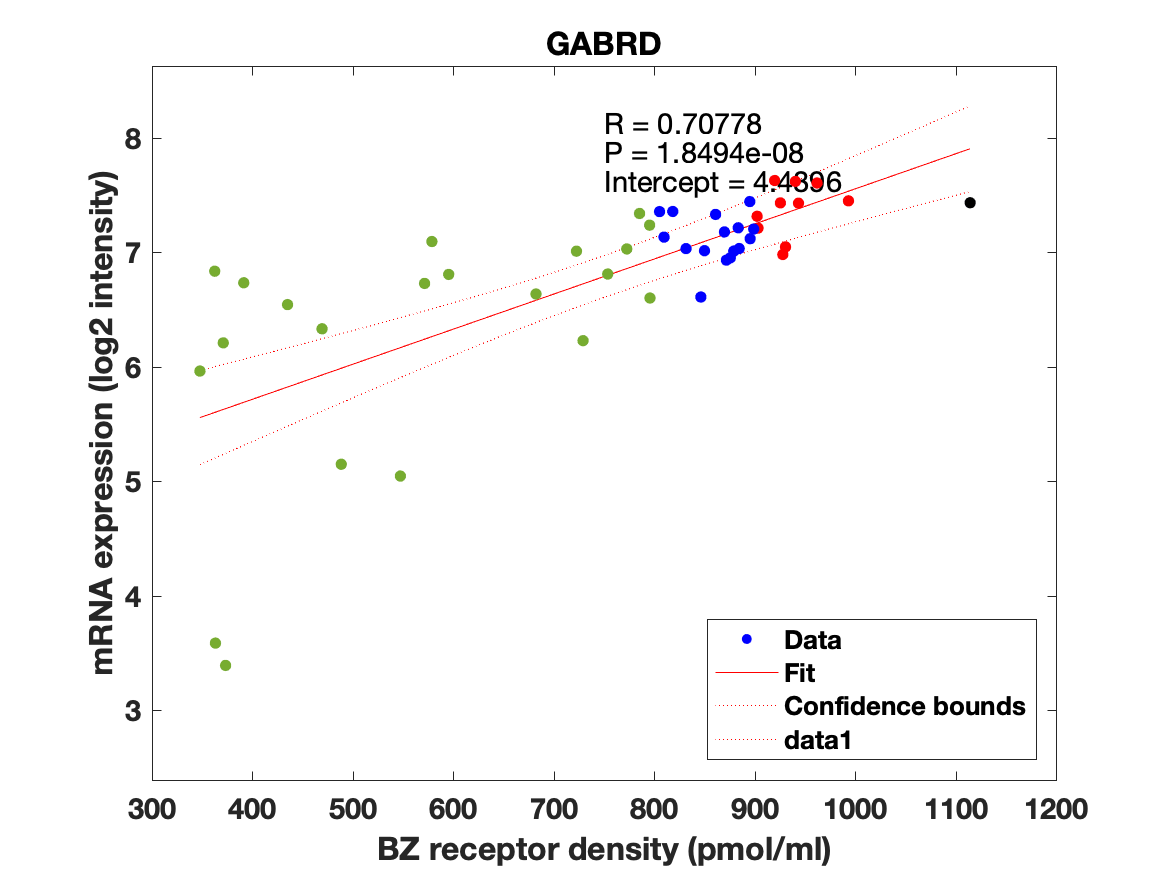

### GABRE.png

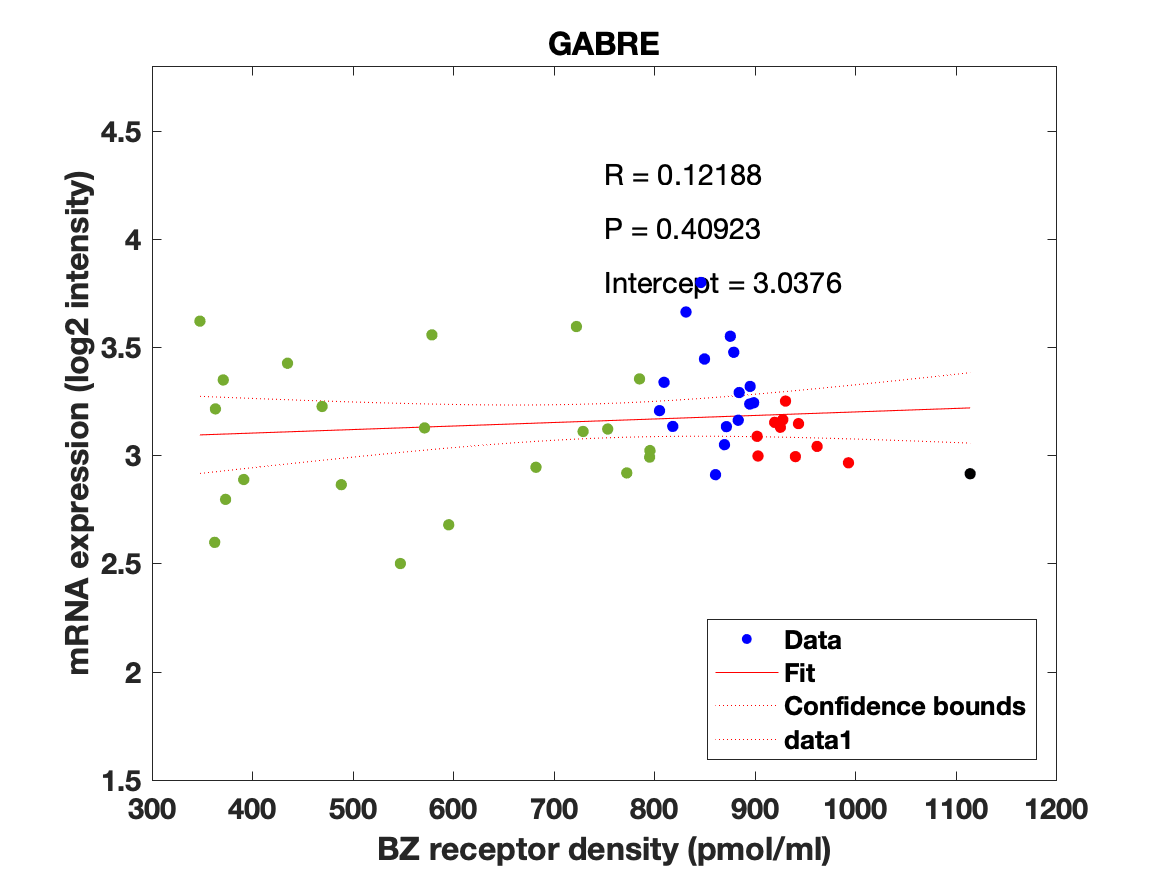

### GABRG1.png

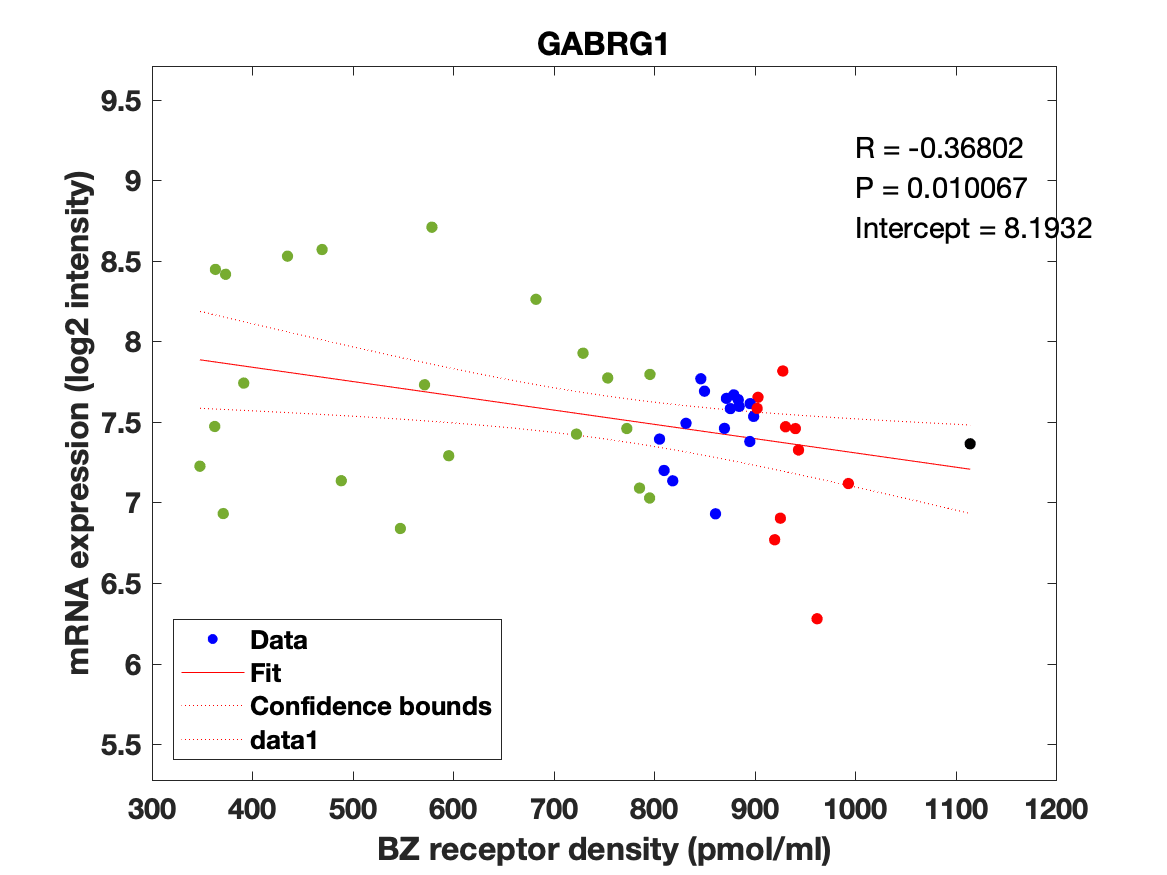

### GABRG2.png

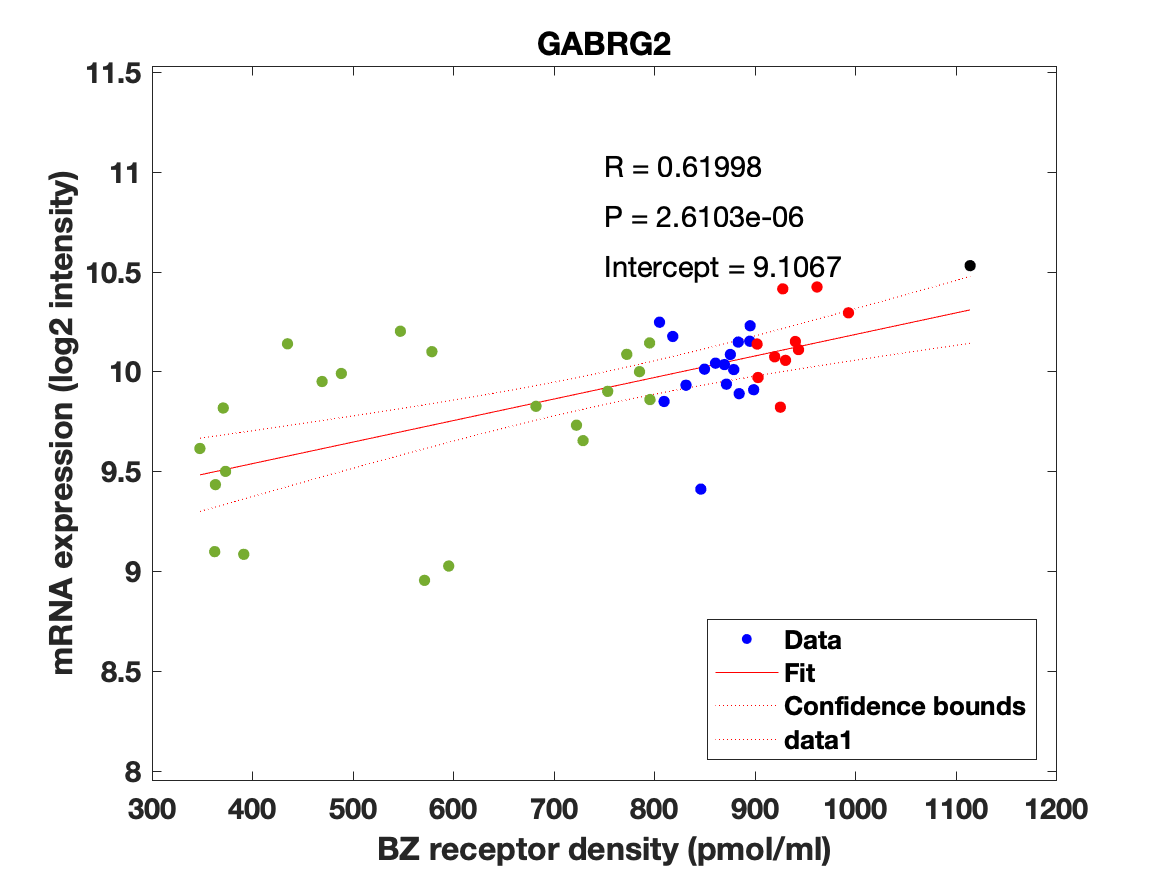

### GABRG3.png

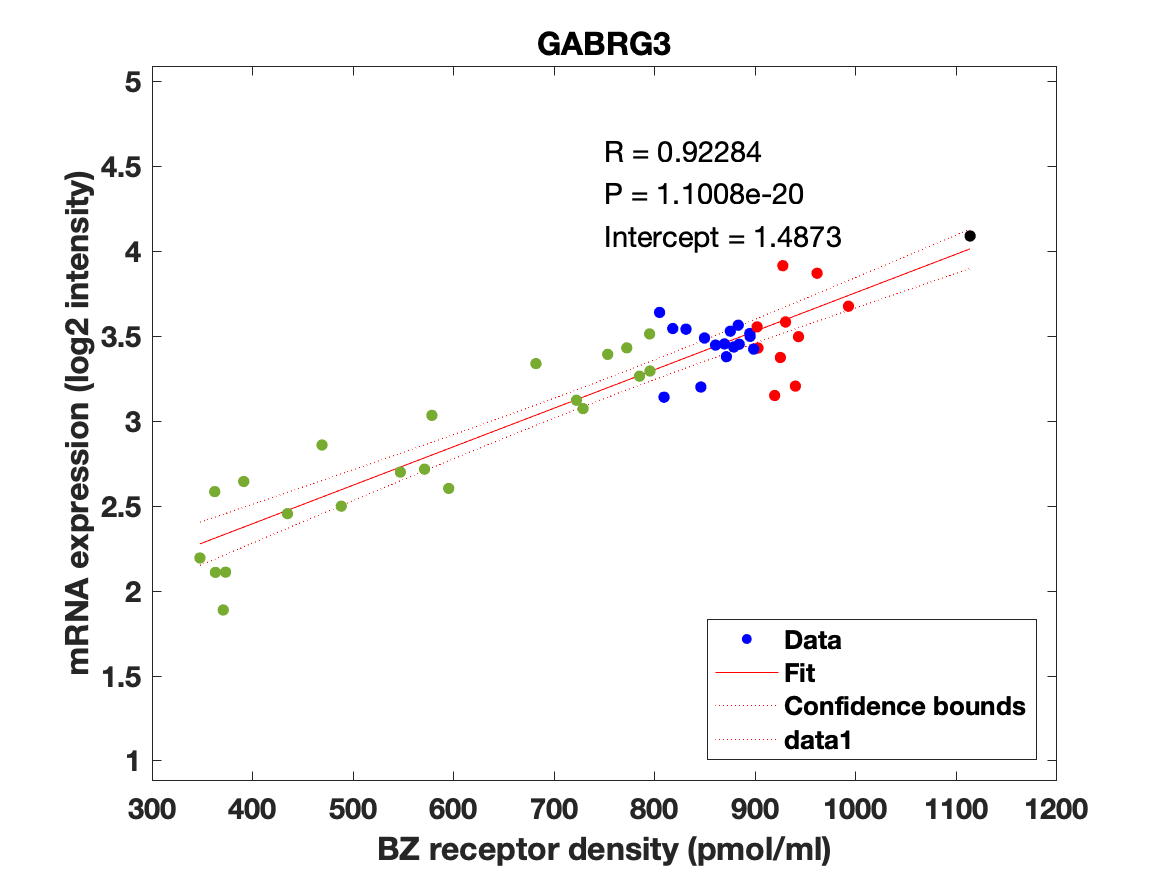

### GABRP.png

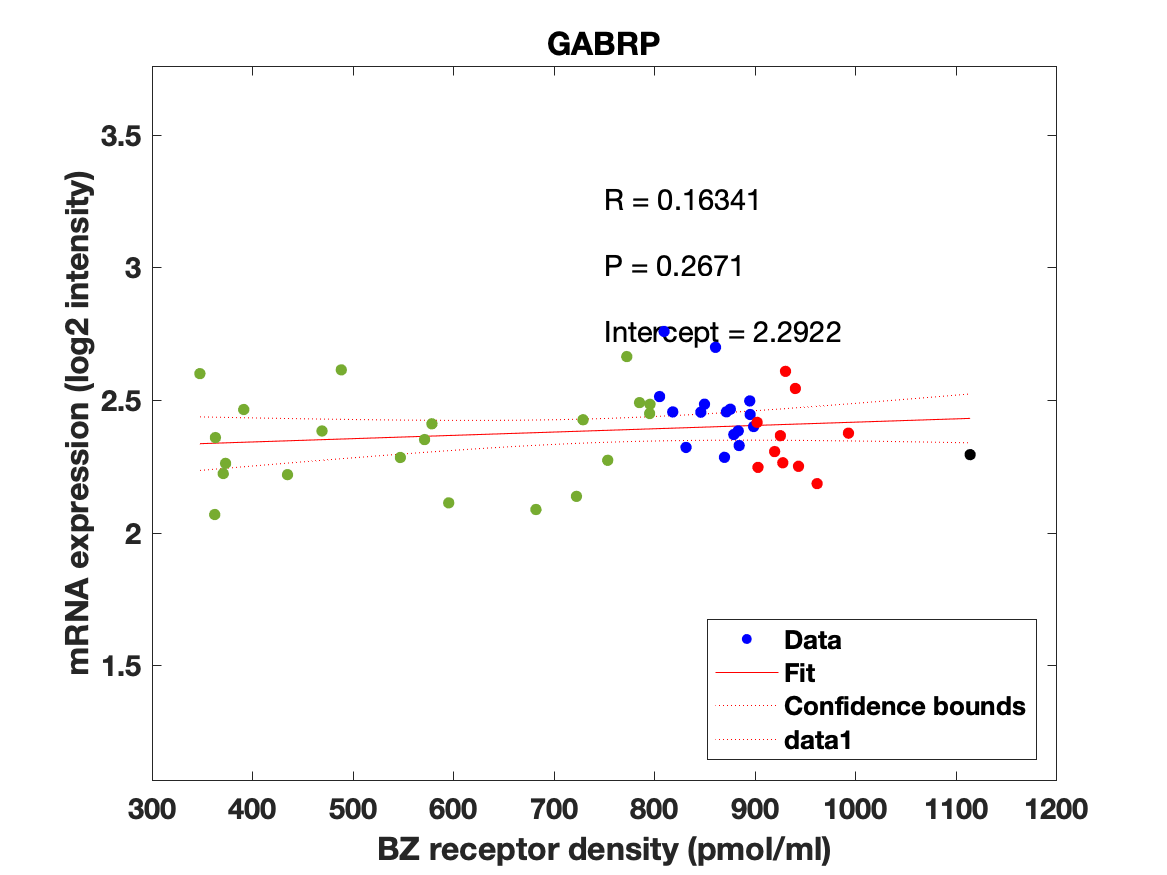

### GABRQ.png

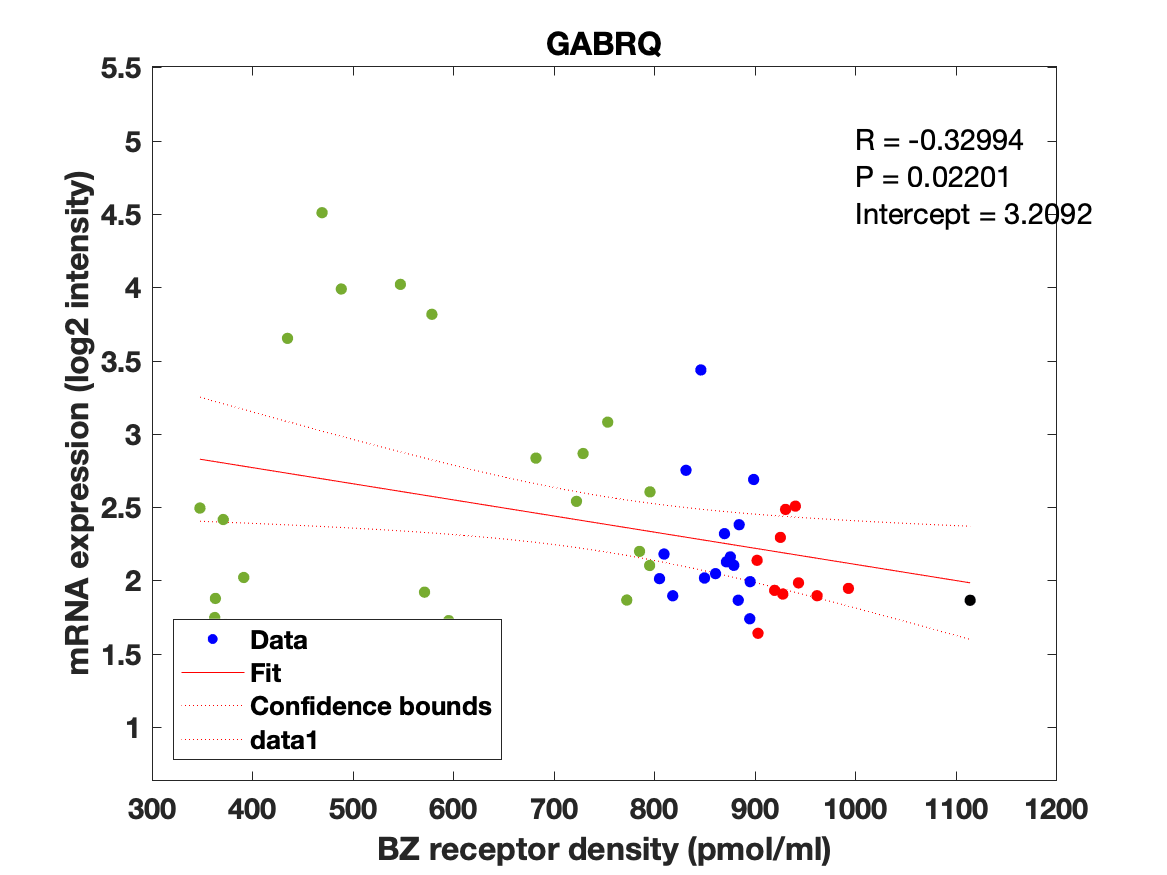

### GABRR1.png

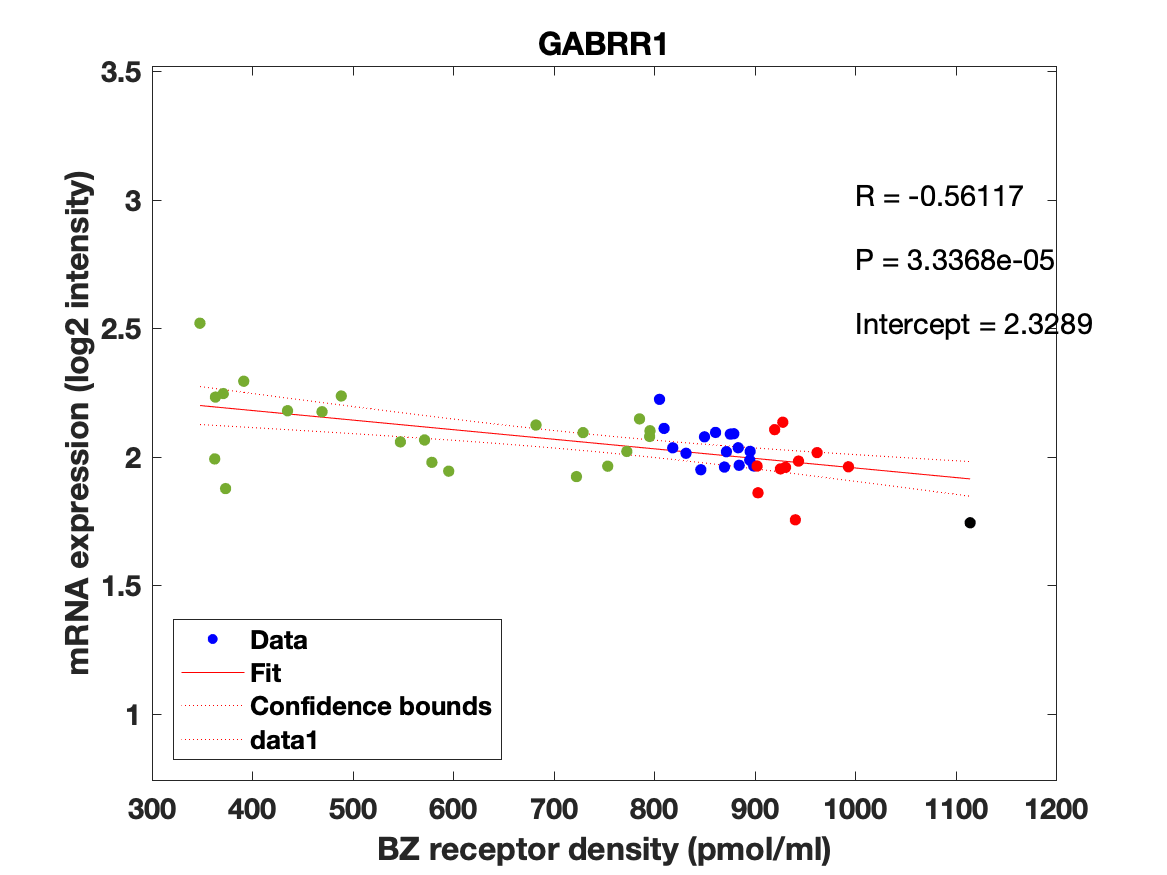

### GABRR2.png

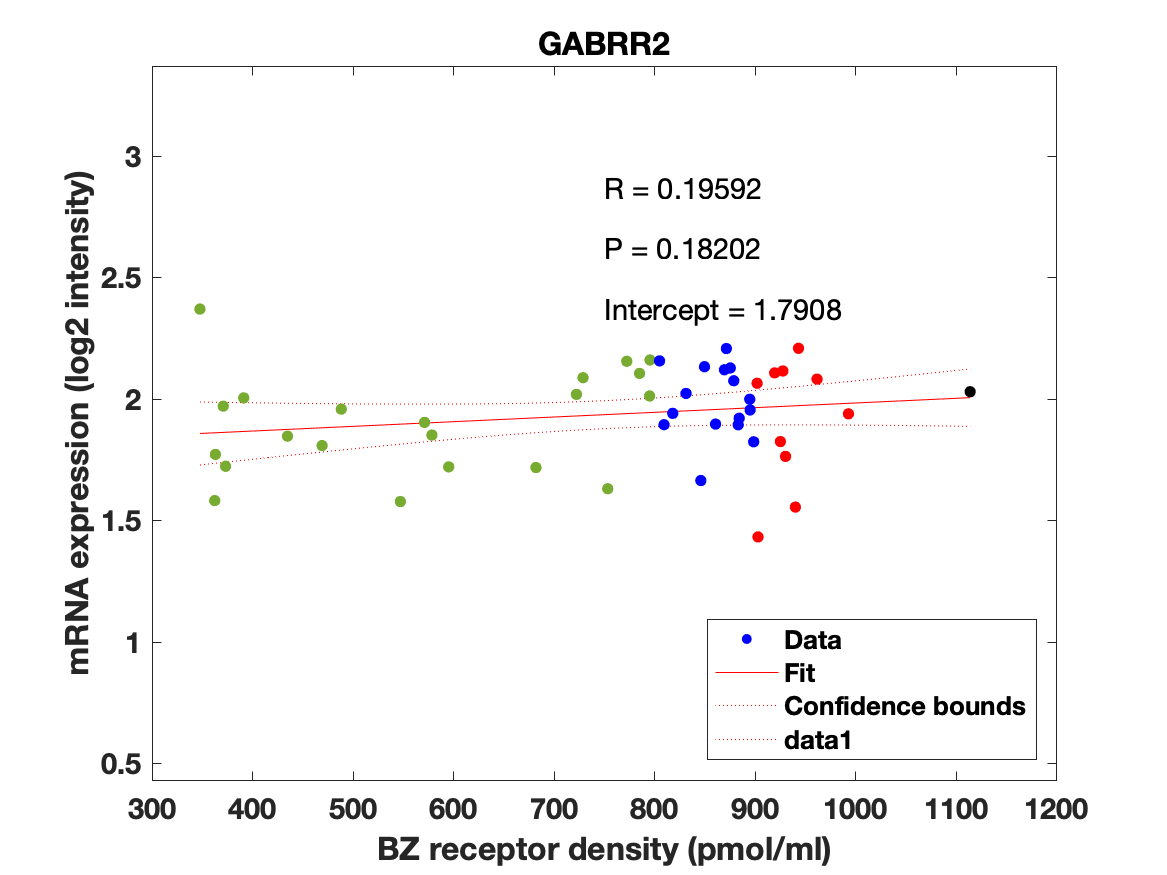

### GABRR3.png

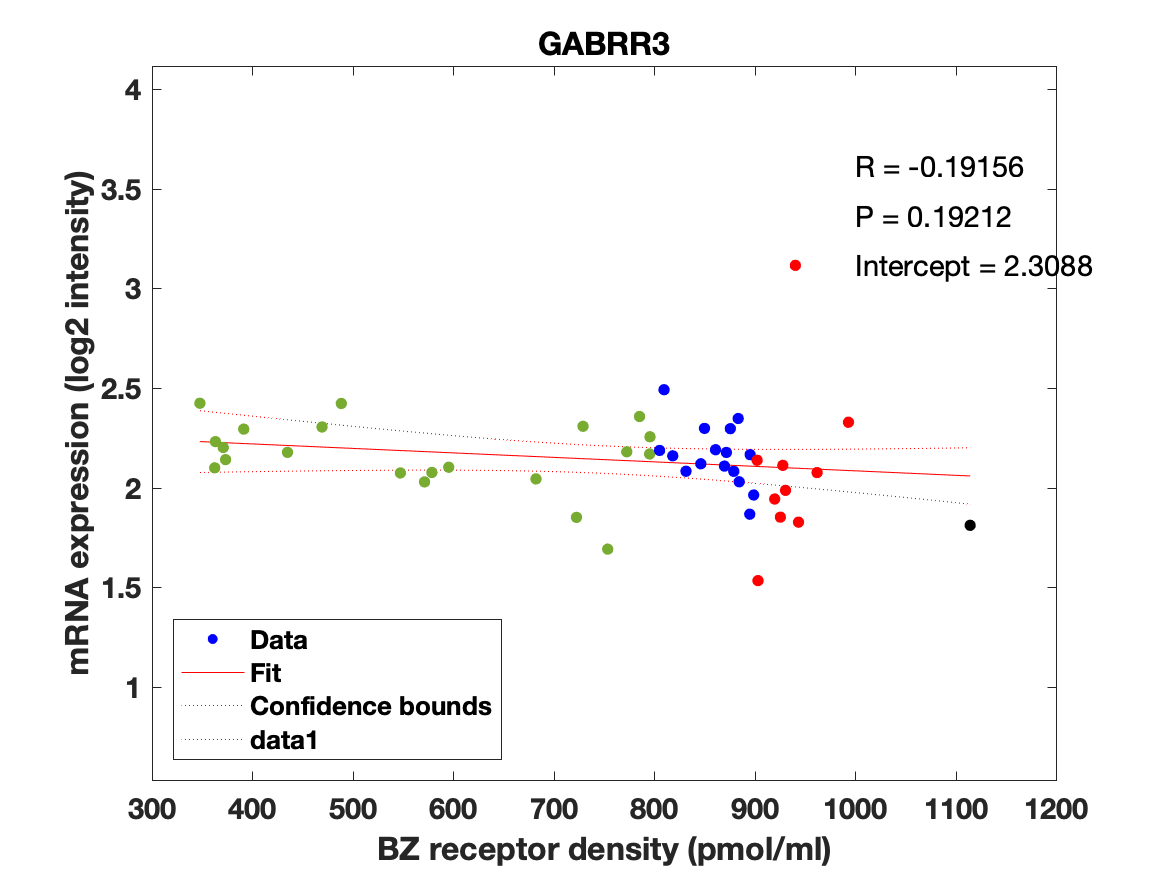

### group_fit_95conf.png

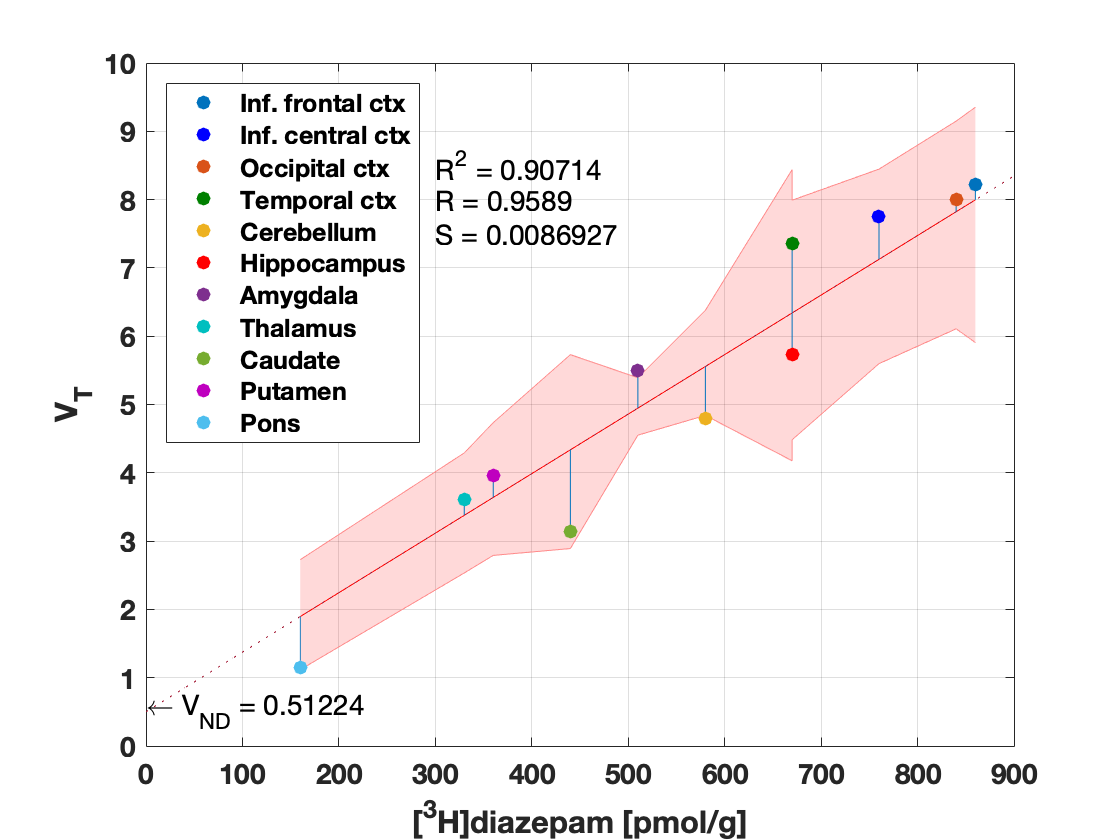

### subj1_s1_fit.png

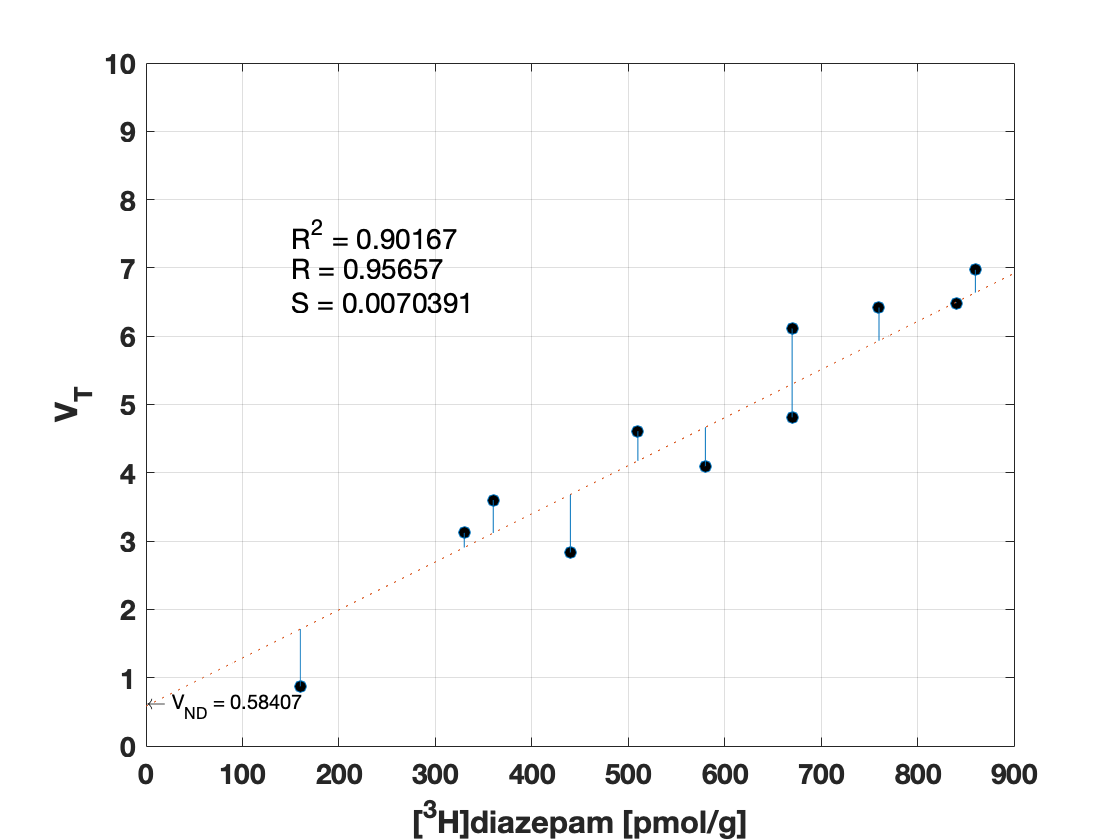

### subj2_s1_fit.png

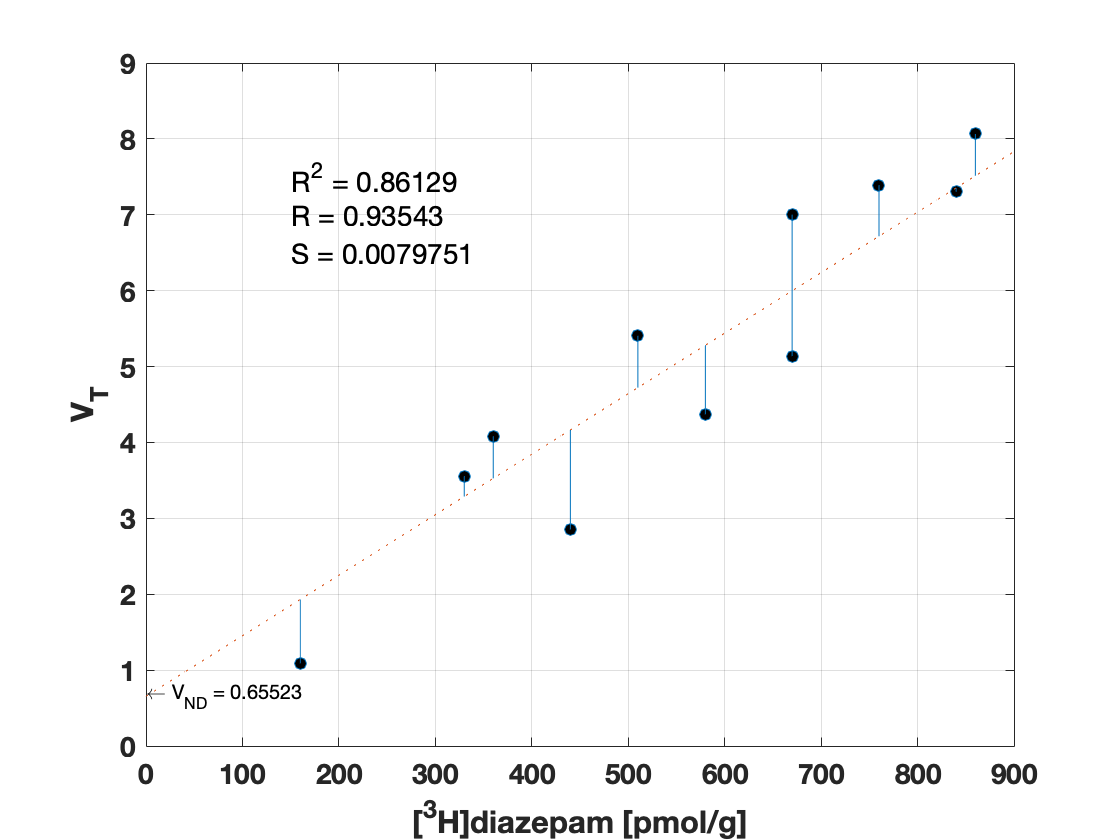

### subj3_s1_fit.png

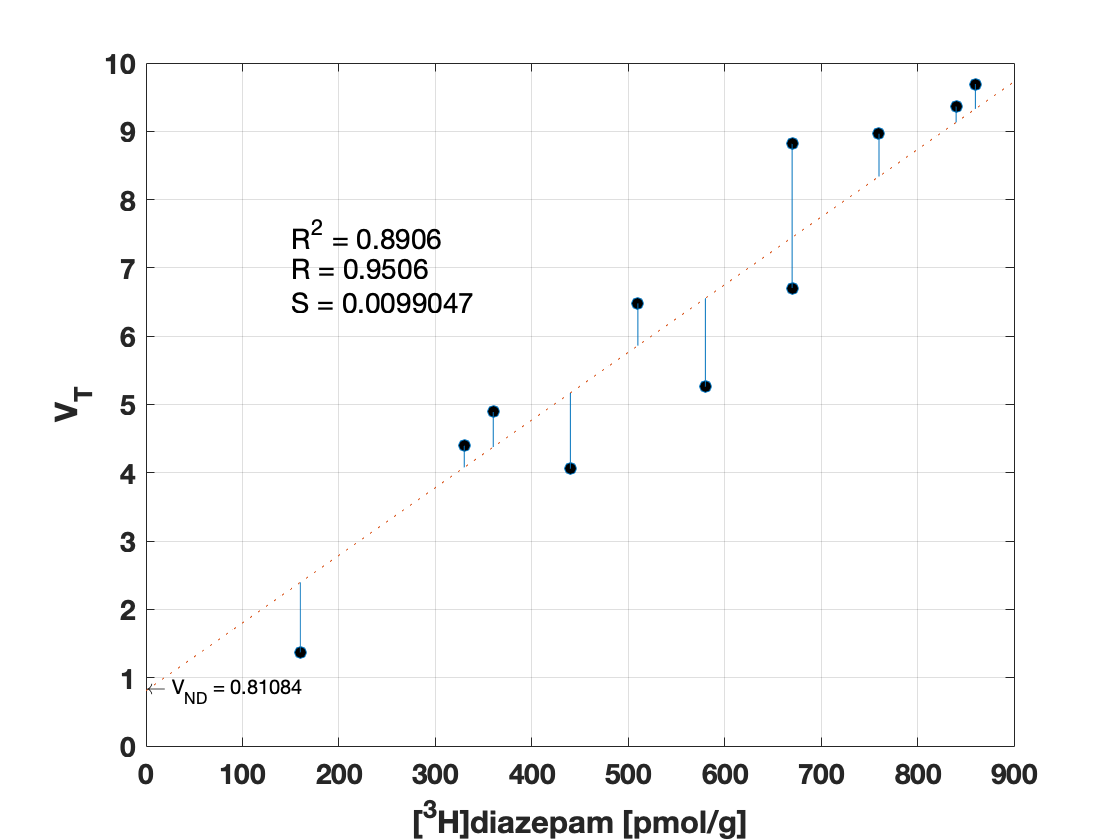

### subj4_s1_fit.png

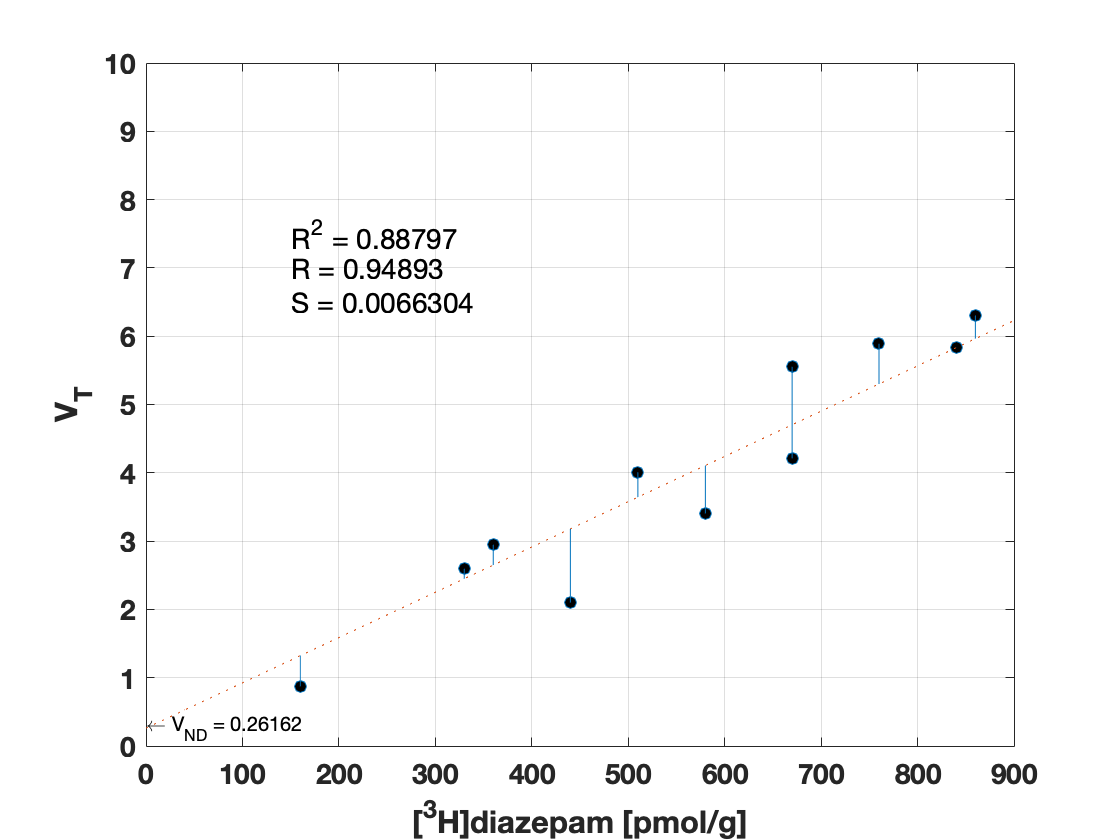

### subj5_s1_fit.png

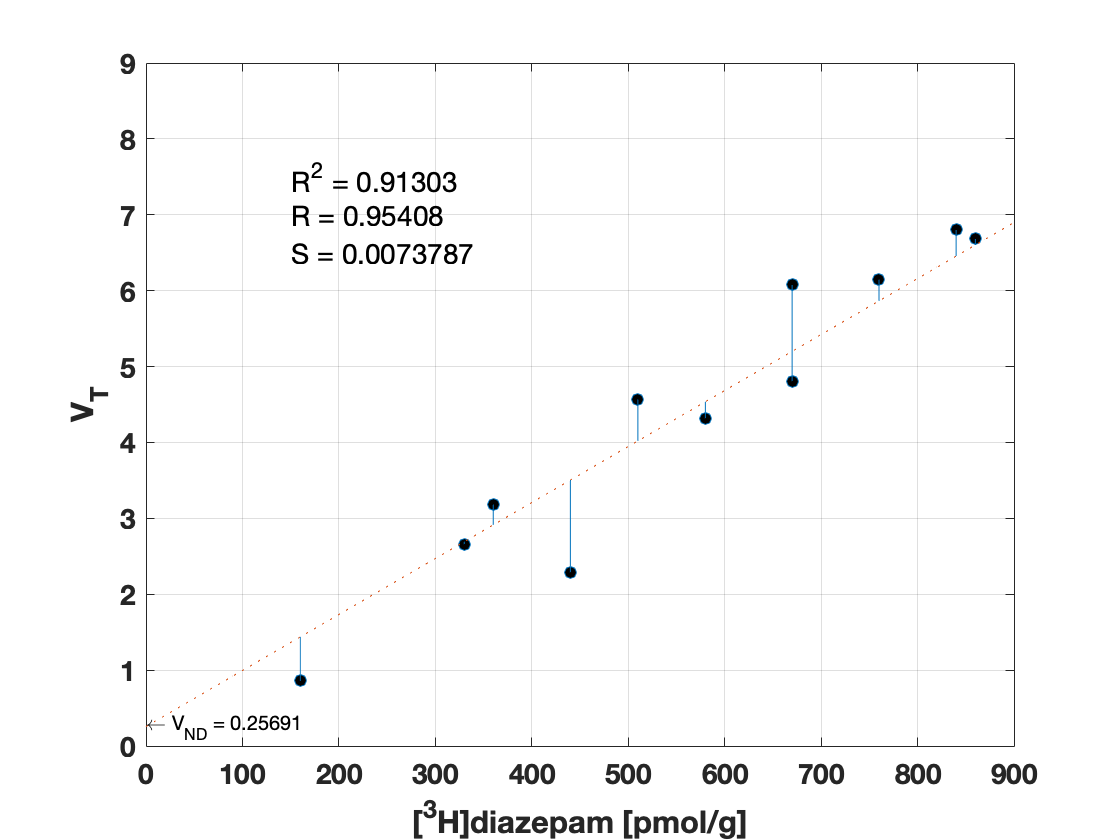

### subj6_s1_fit.png

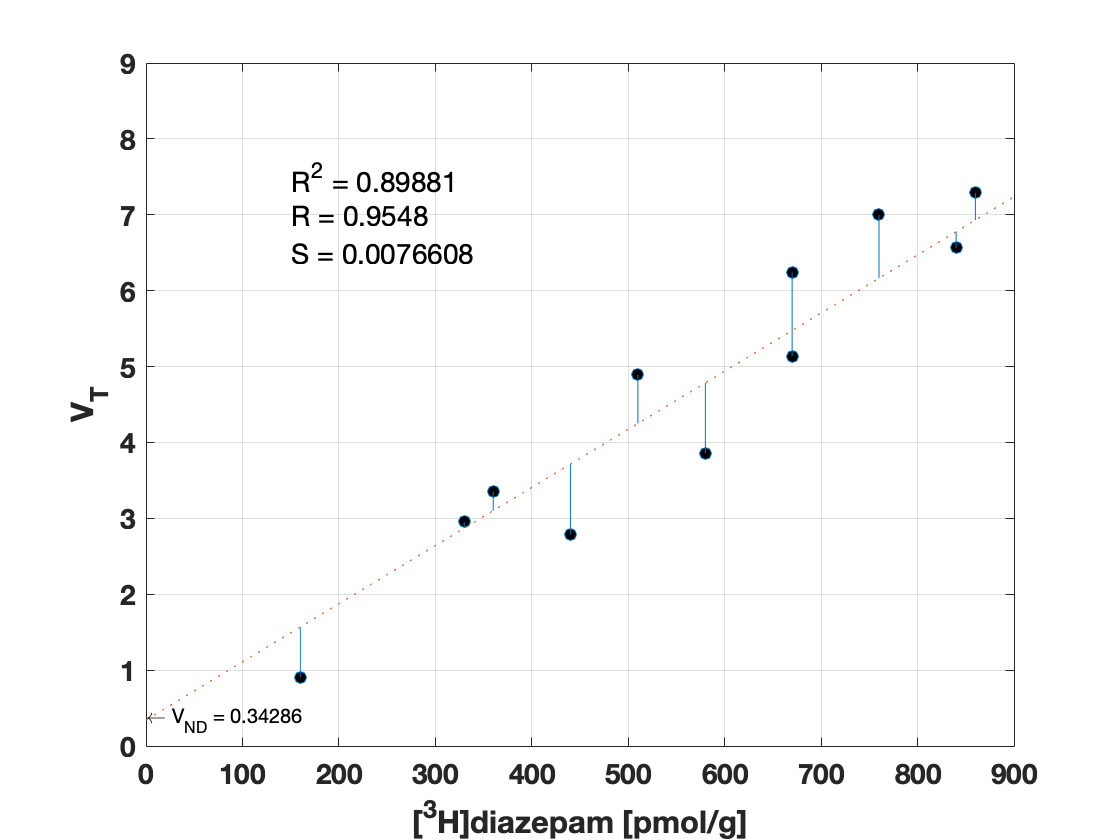

### subj7_s1_fit.png

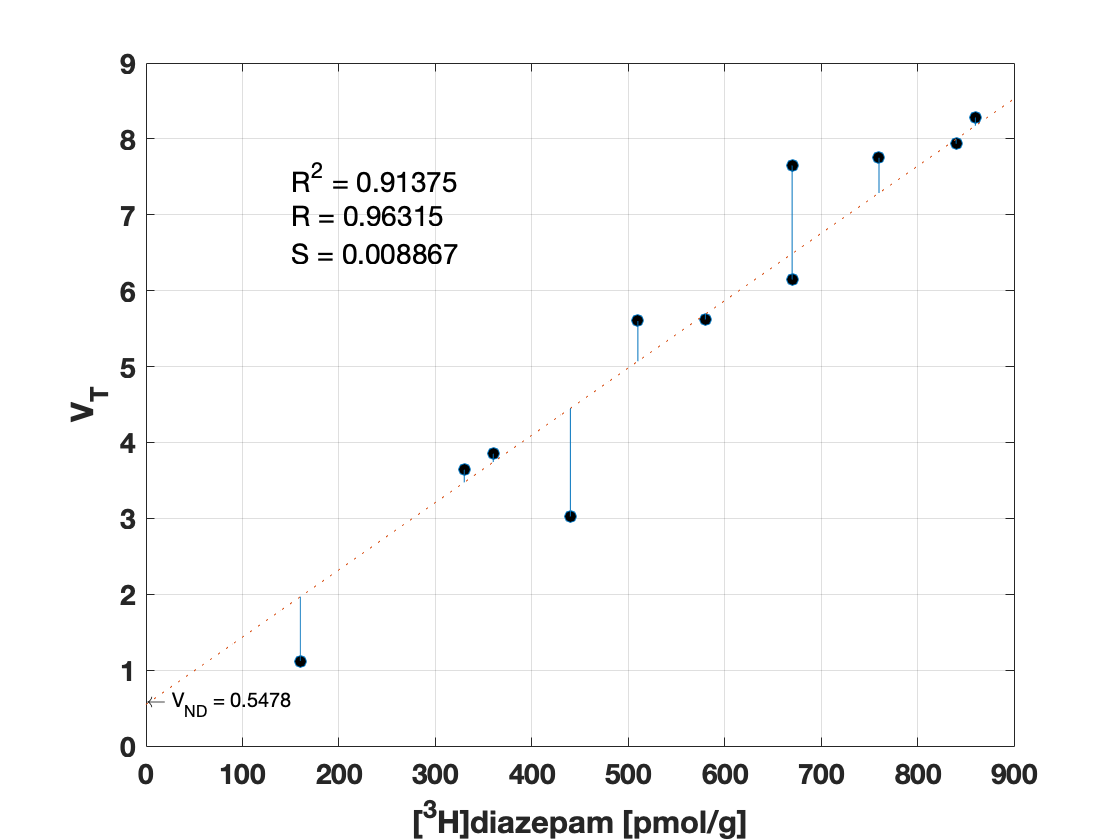

### subj8_s1_fit.png

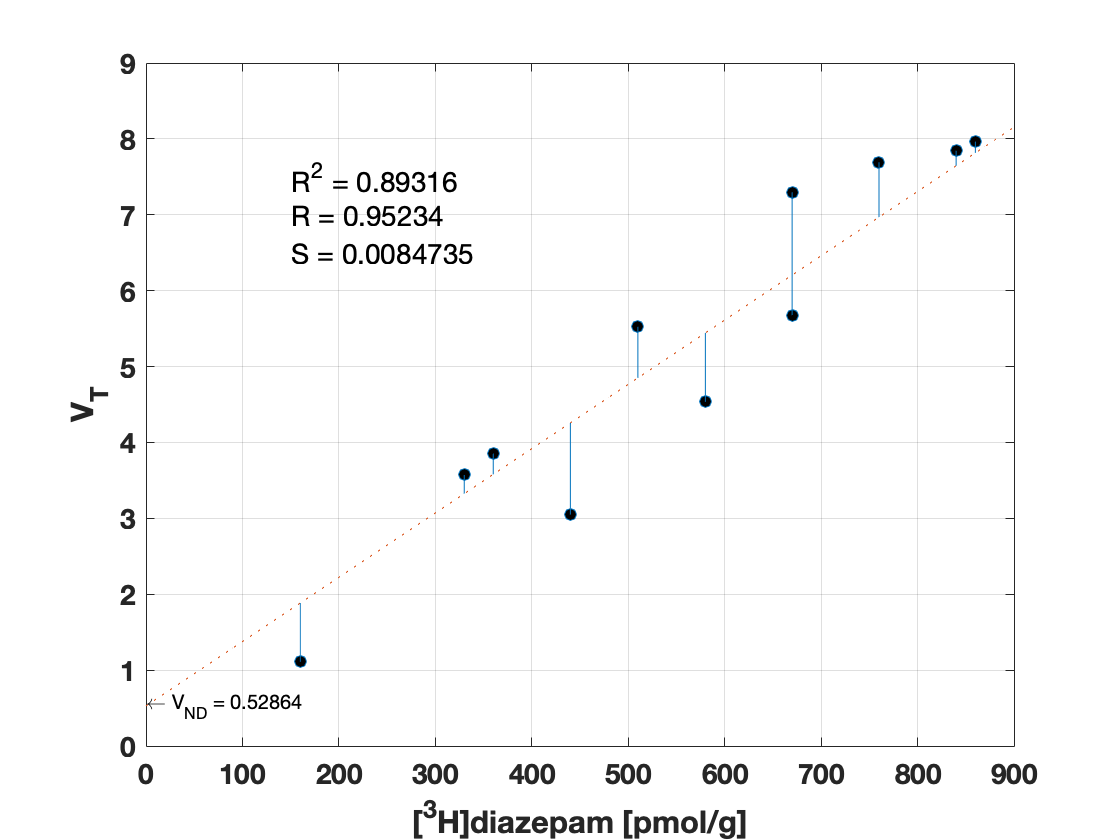

### subj8_s2_fit.png

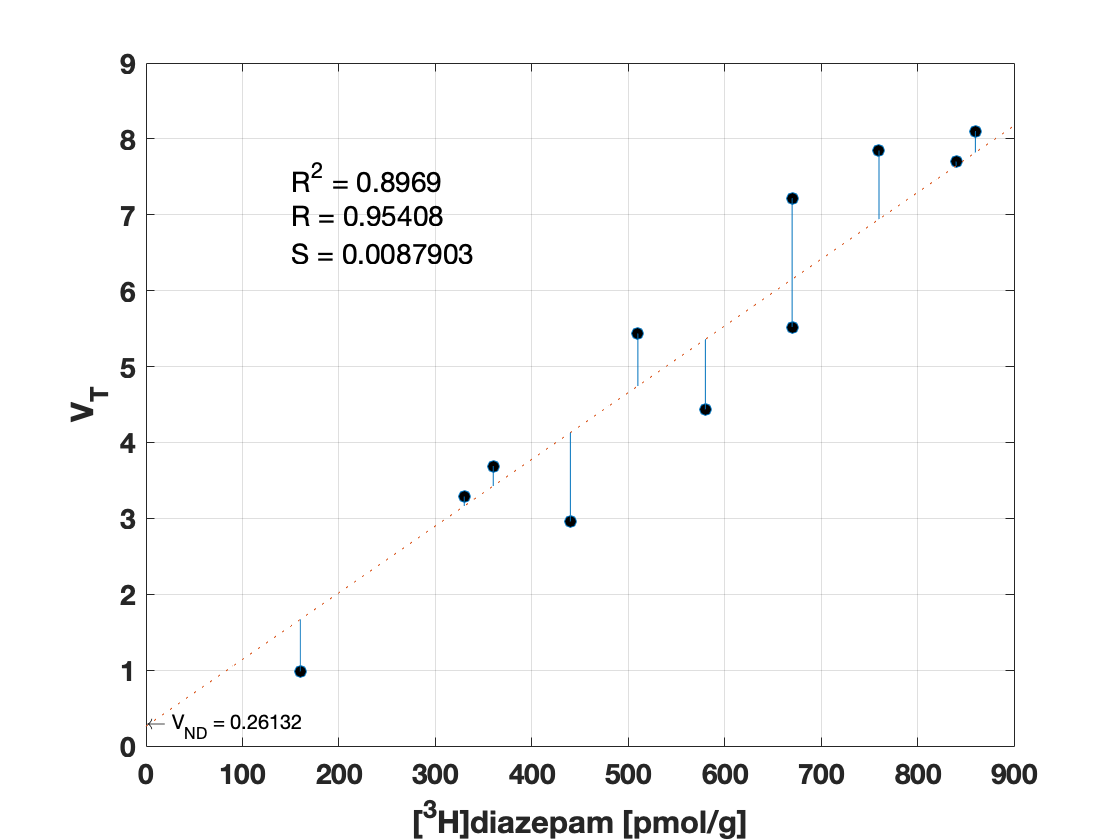

### subj9_s1_fit.png

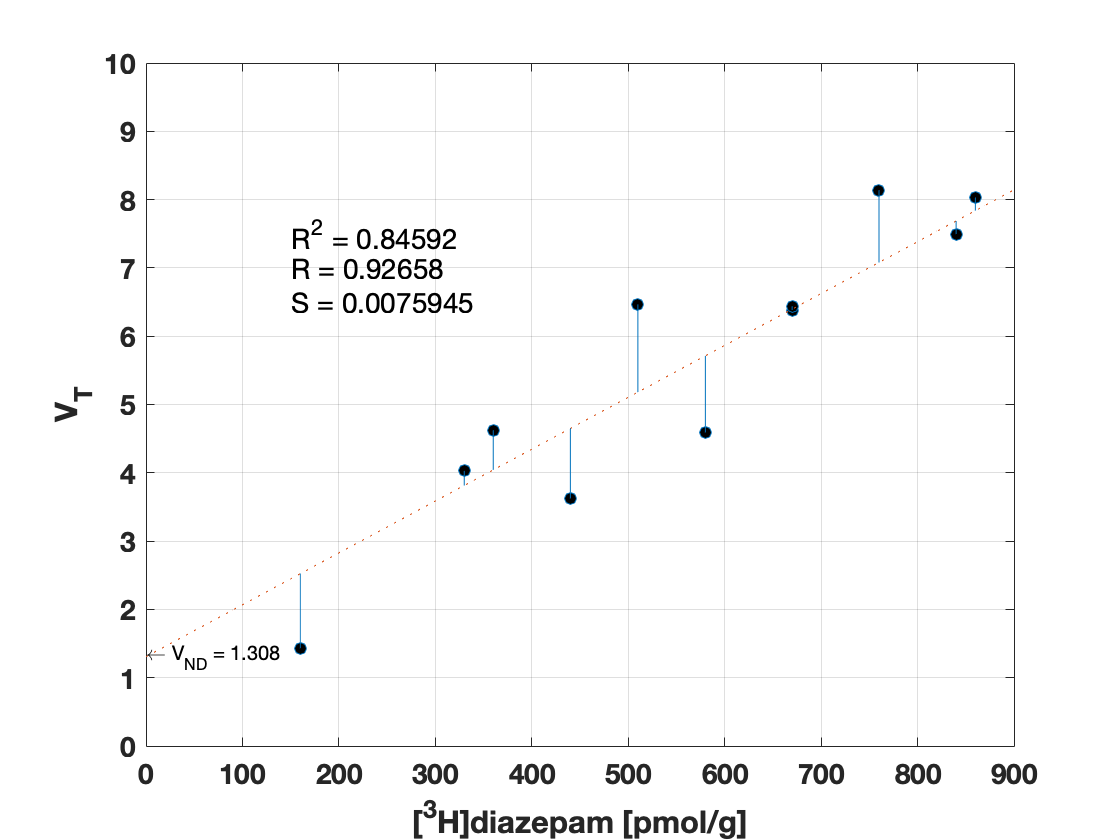

### subj9_s2_fit.png

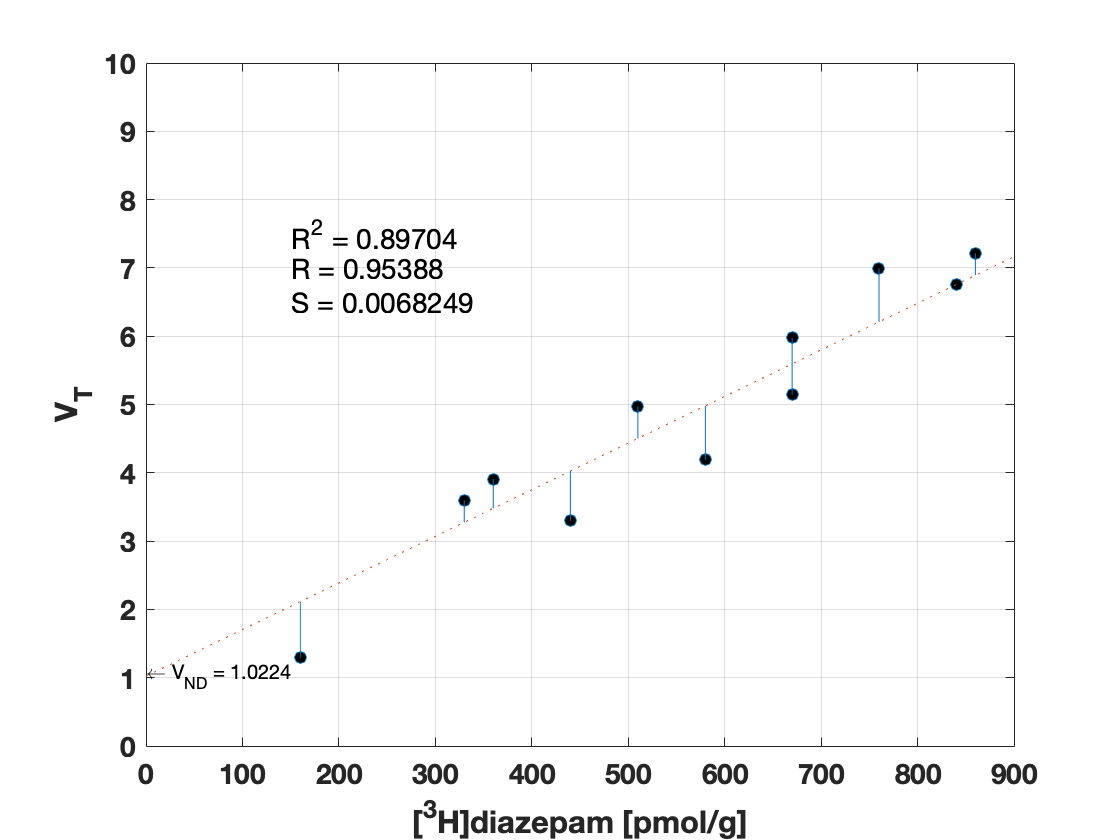

### subj10_s1_fit.png

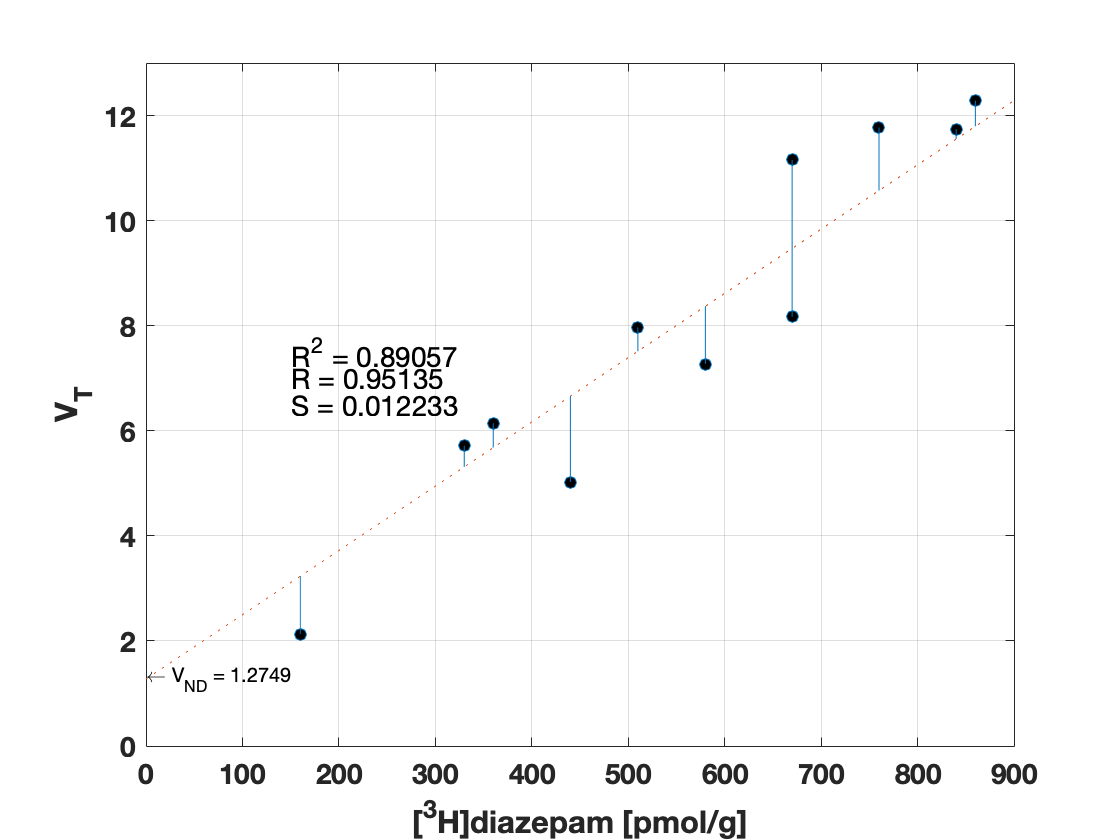

### subj10_s2_fit.png

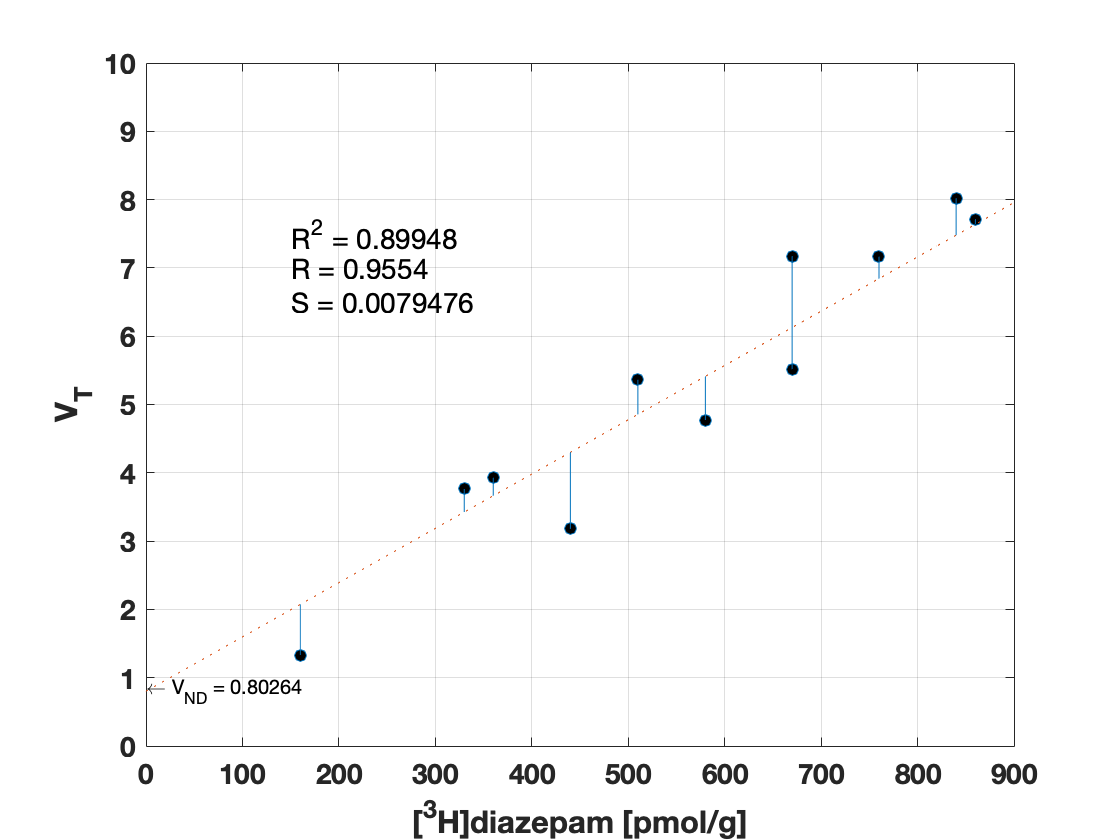

### subj11_s1_fit.png

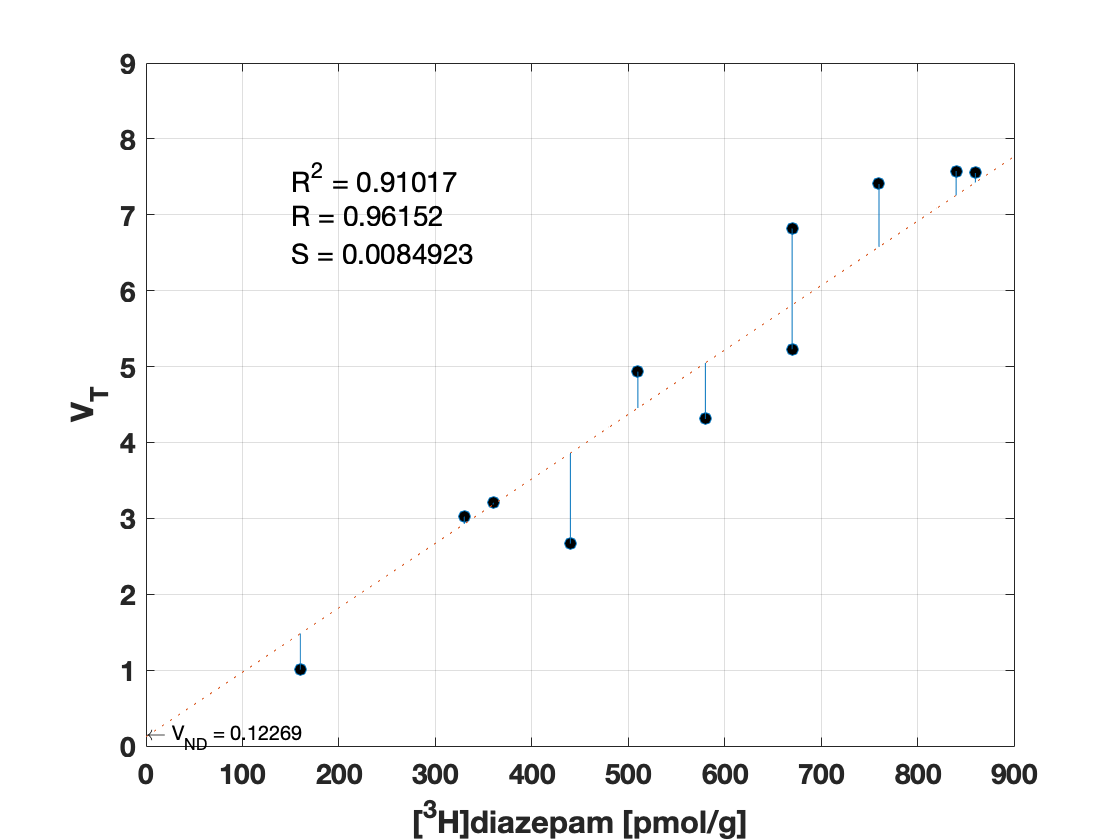

### subj11_s2_fit.png

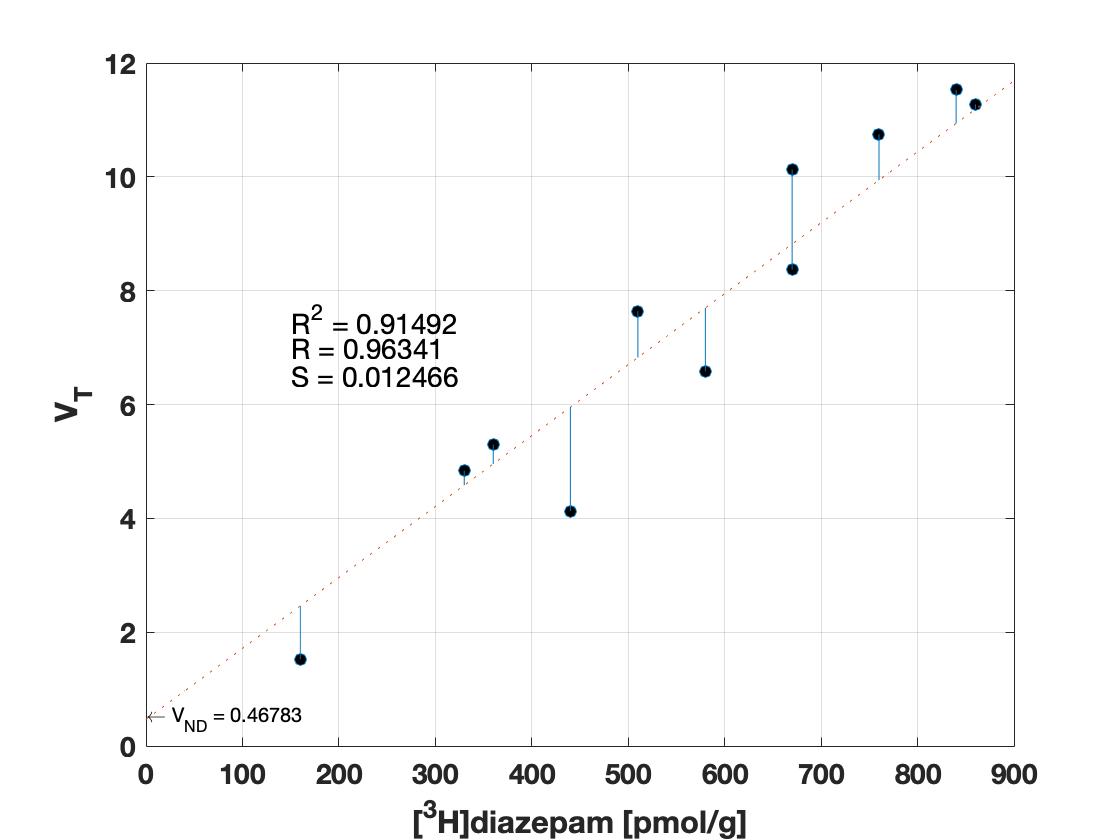

### subj12_s1_fit.png

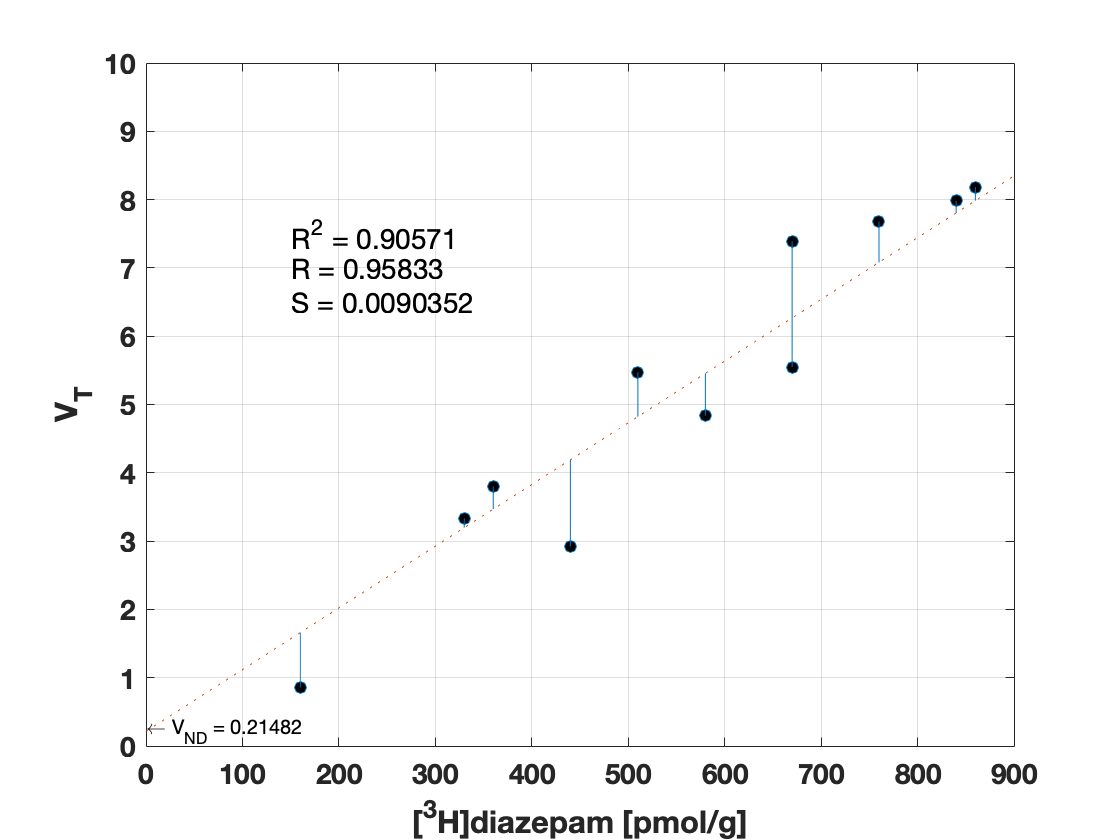

### subj12_s2_fit.png

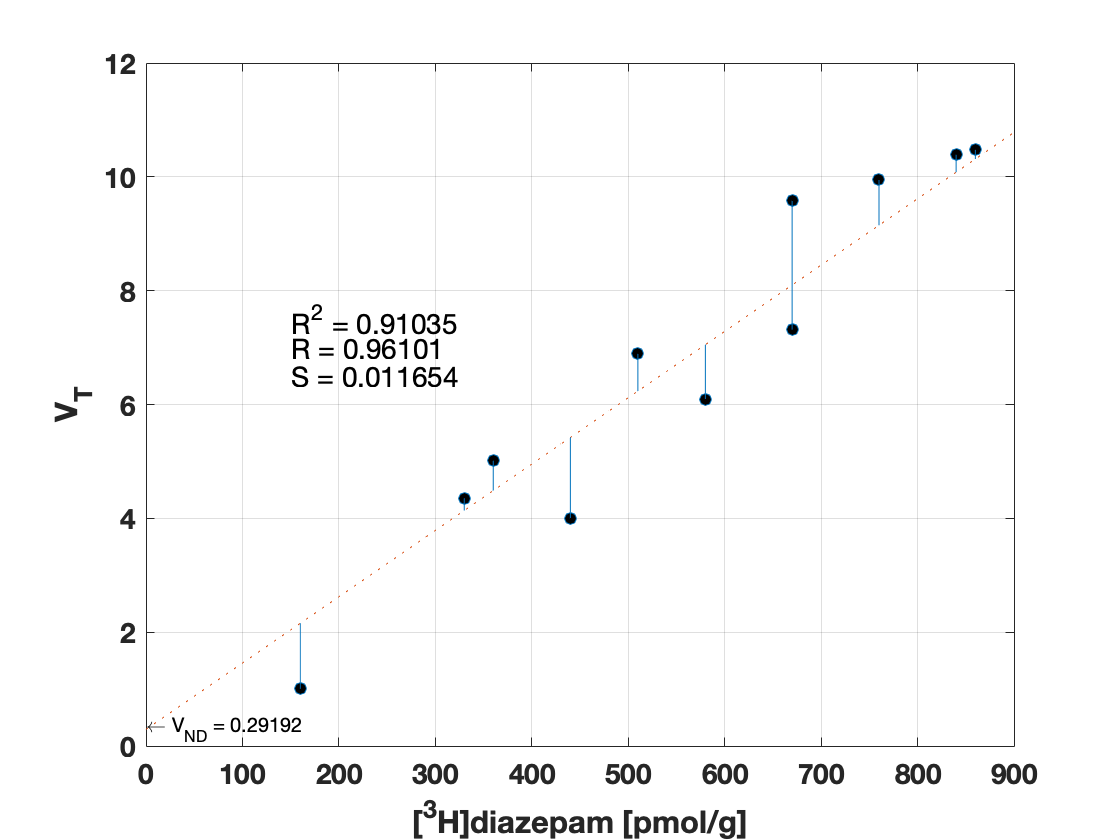

### subj13_s1_fit.png

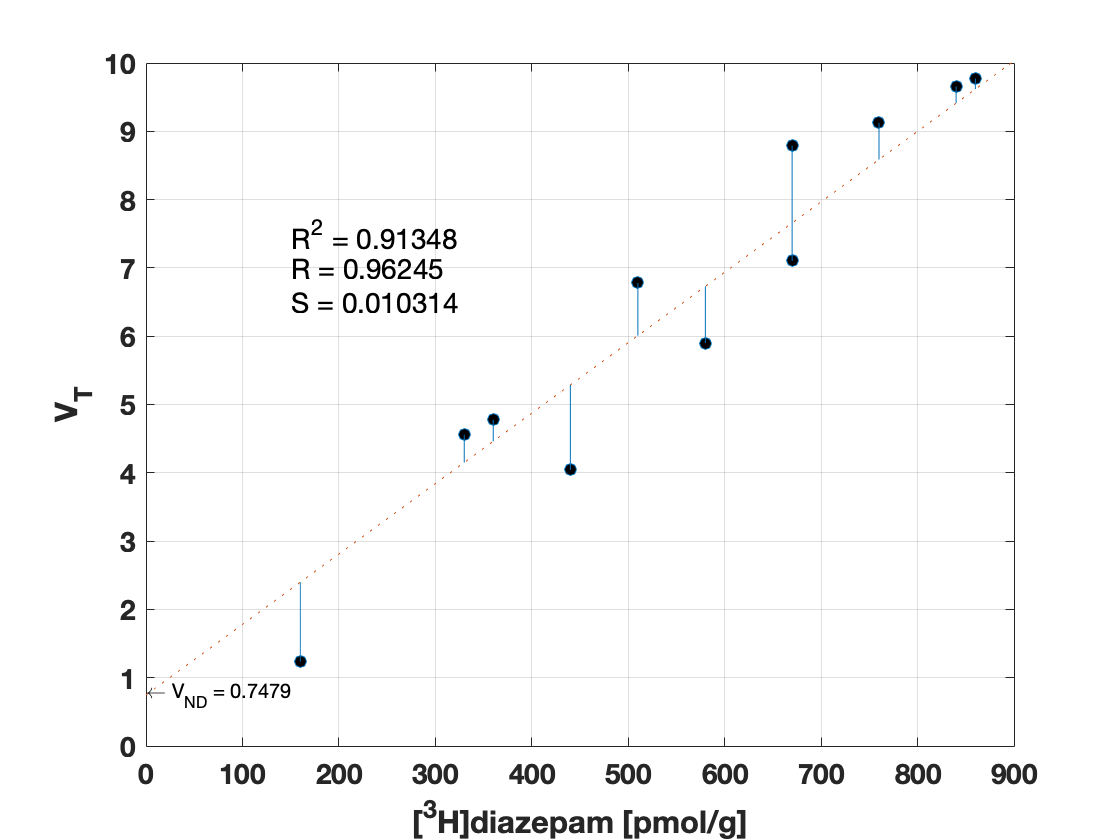

### subj13_s2_fit.png

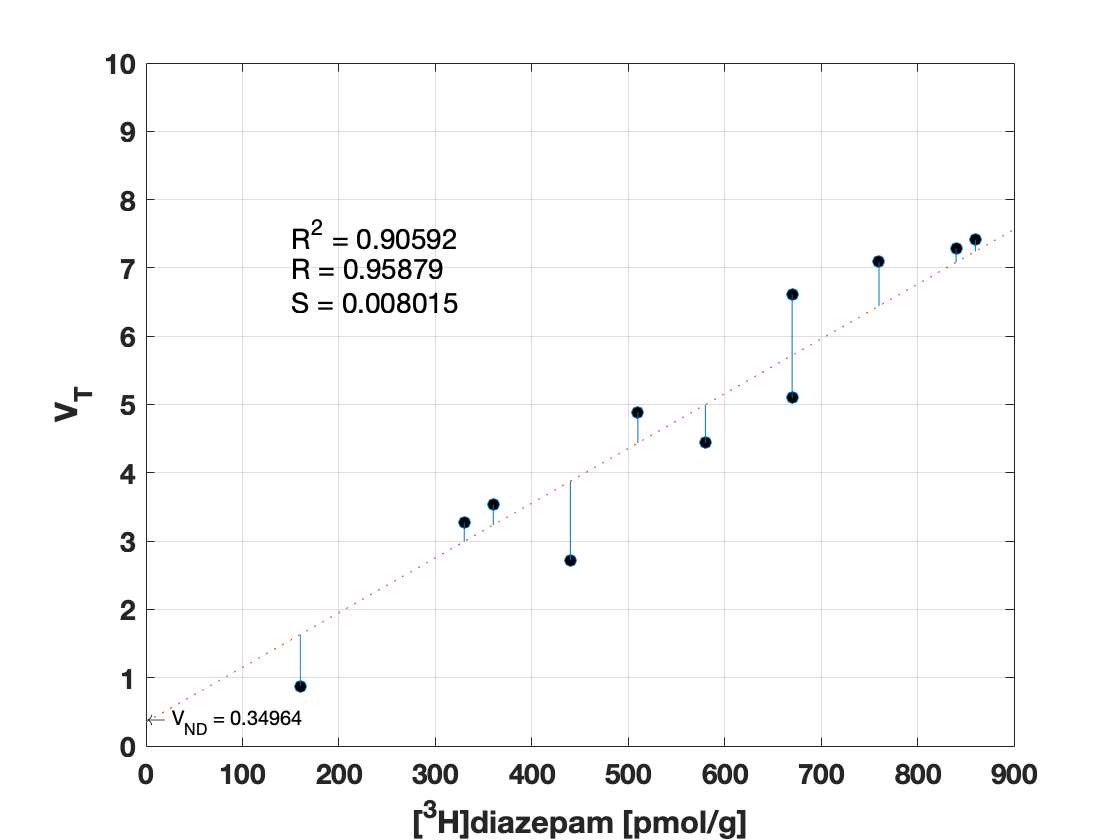

### subj14_s1_fit.png

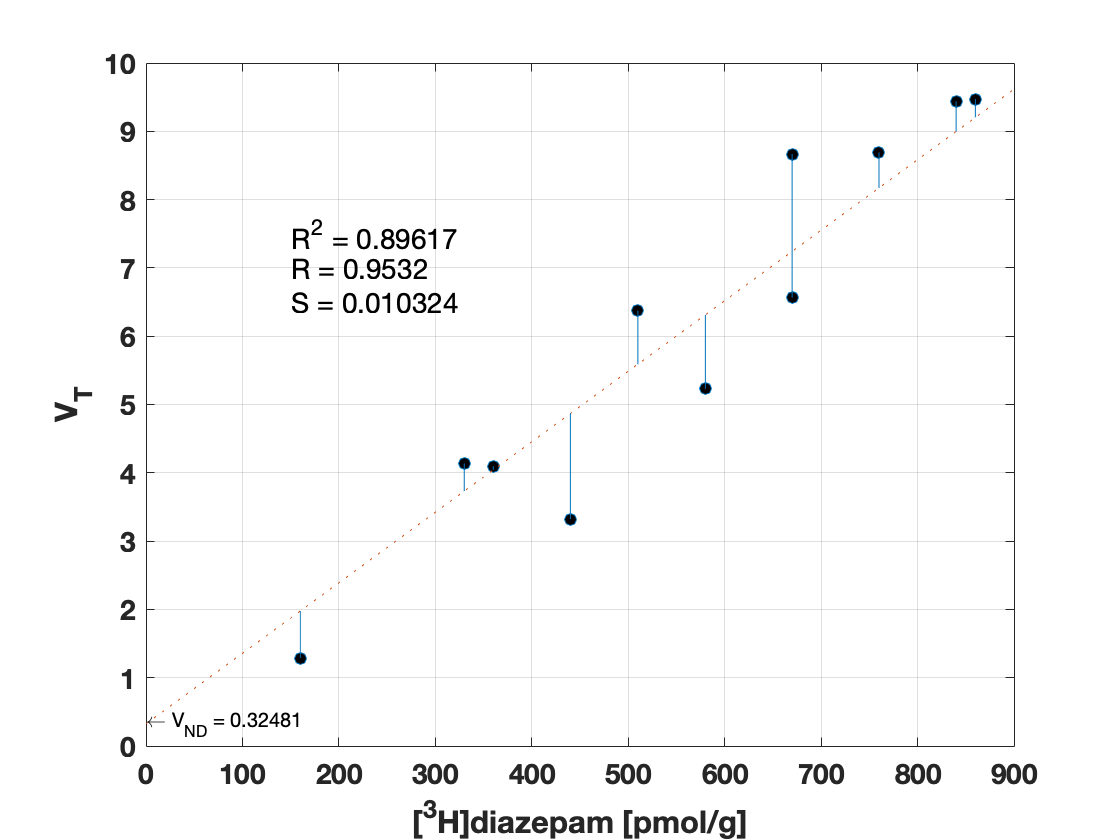

### subj14_s2_fit.png

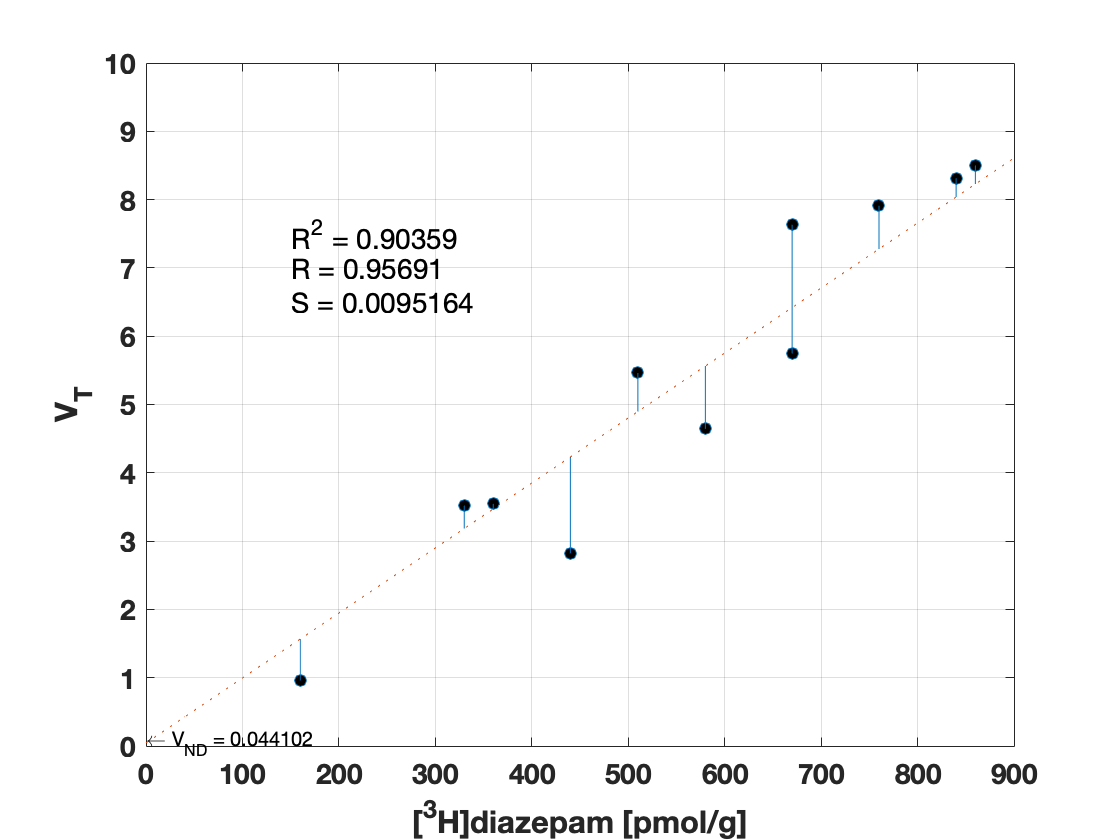

### subj15_s1_fit.png

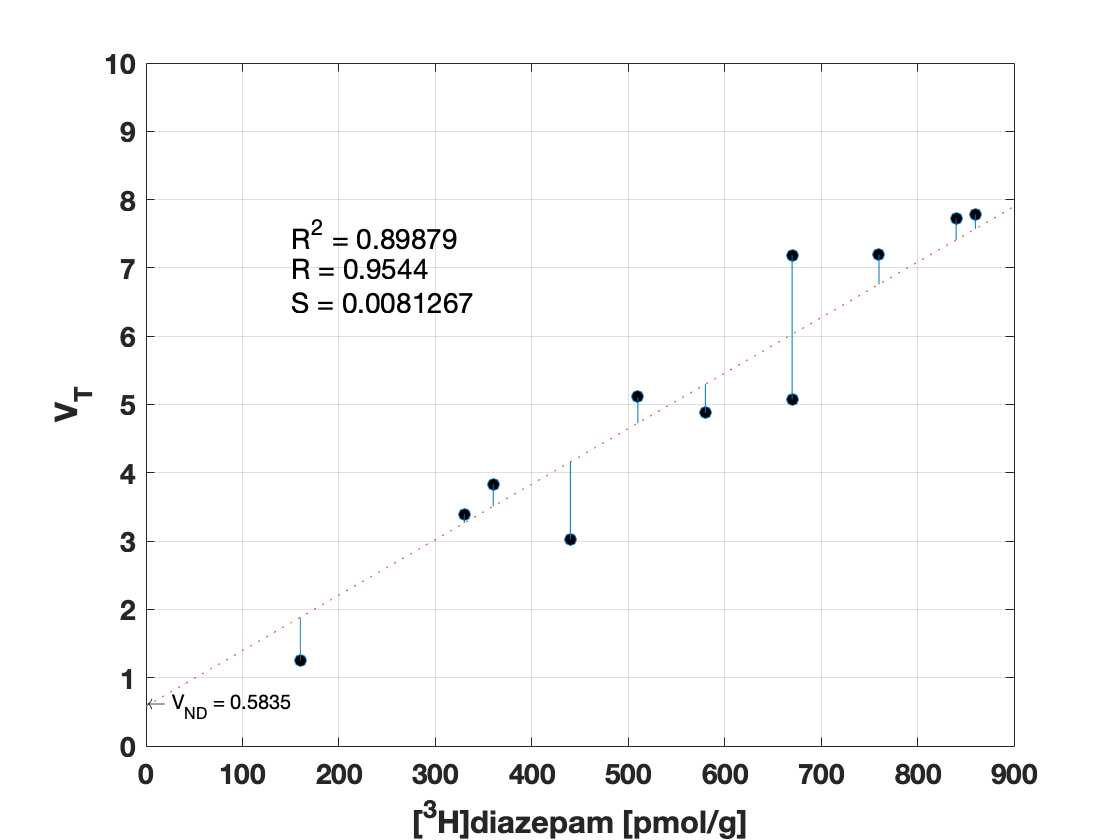

### subj15_s2_fit.png

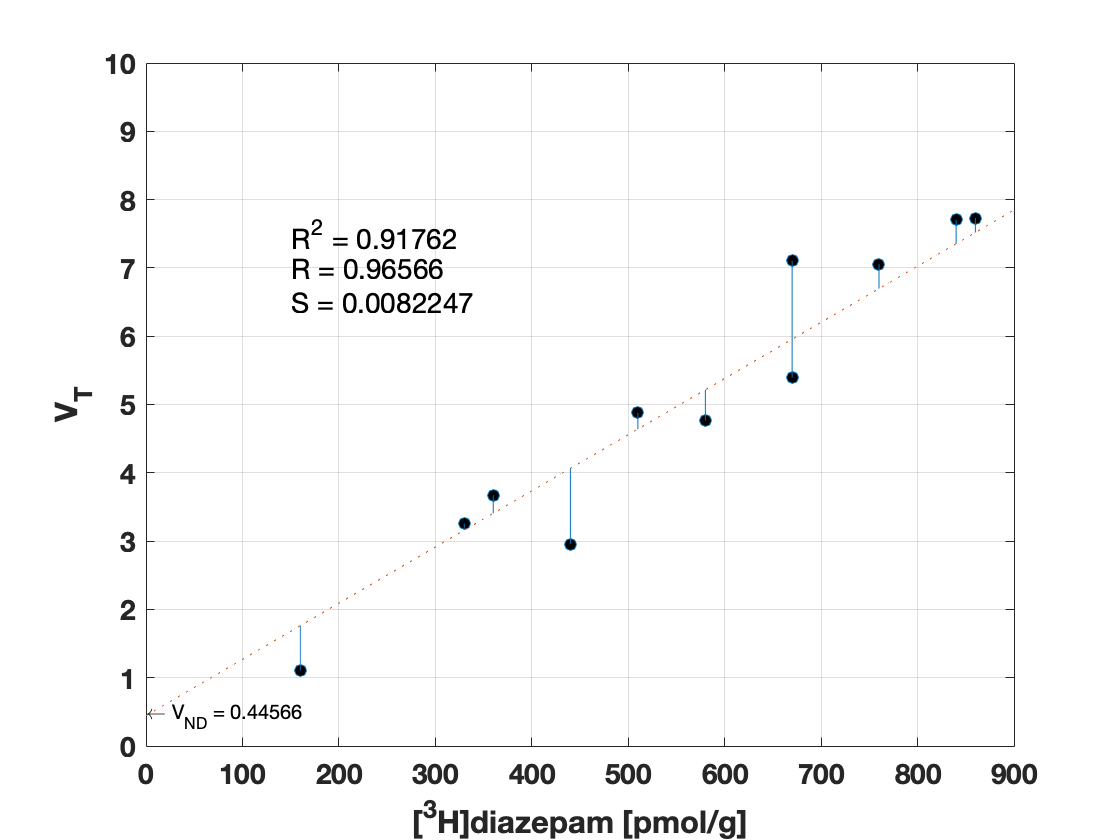

### subj15_s3_fit.png

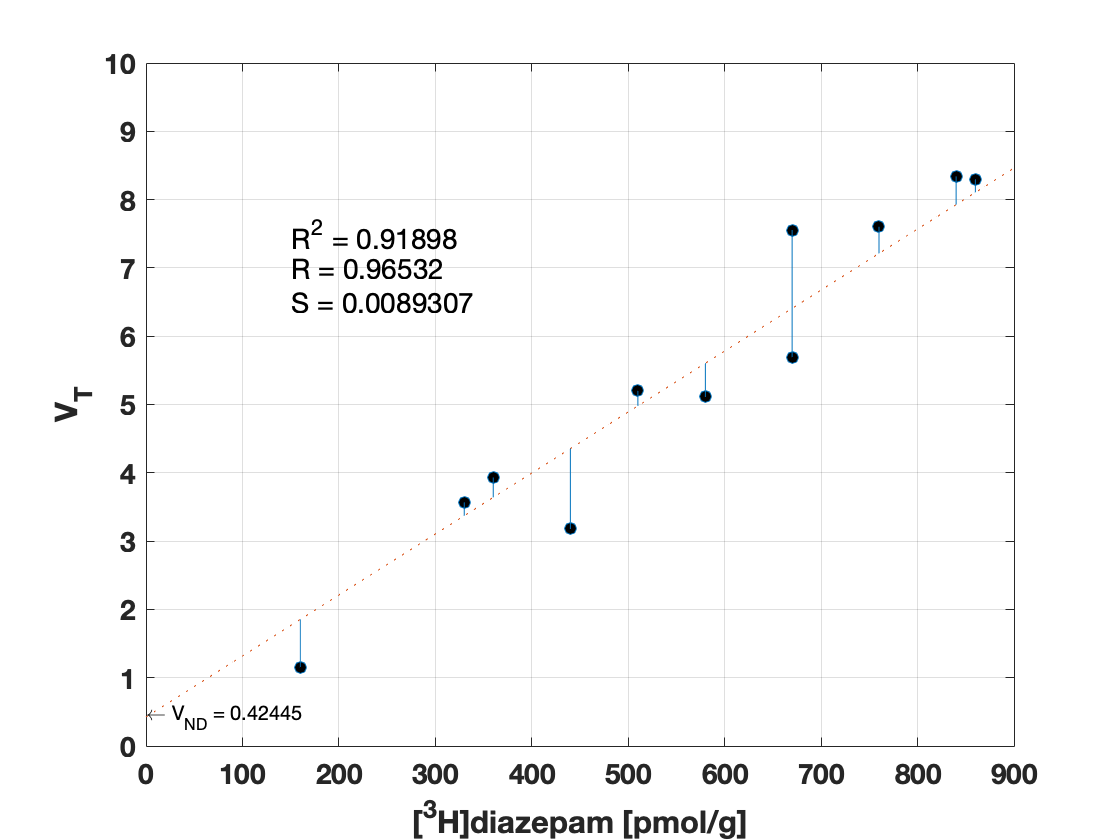

### subj16_s1_fit.png

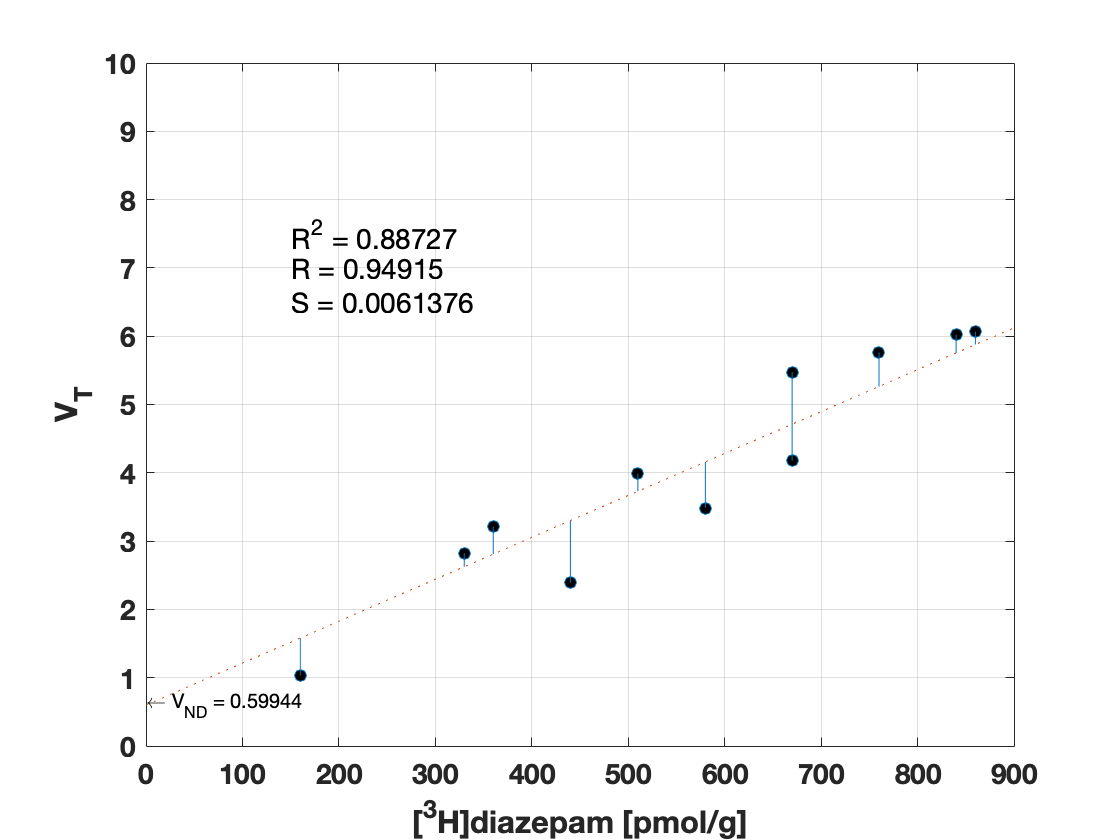

### subj16_s2_fit.png

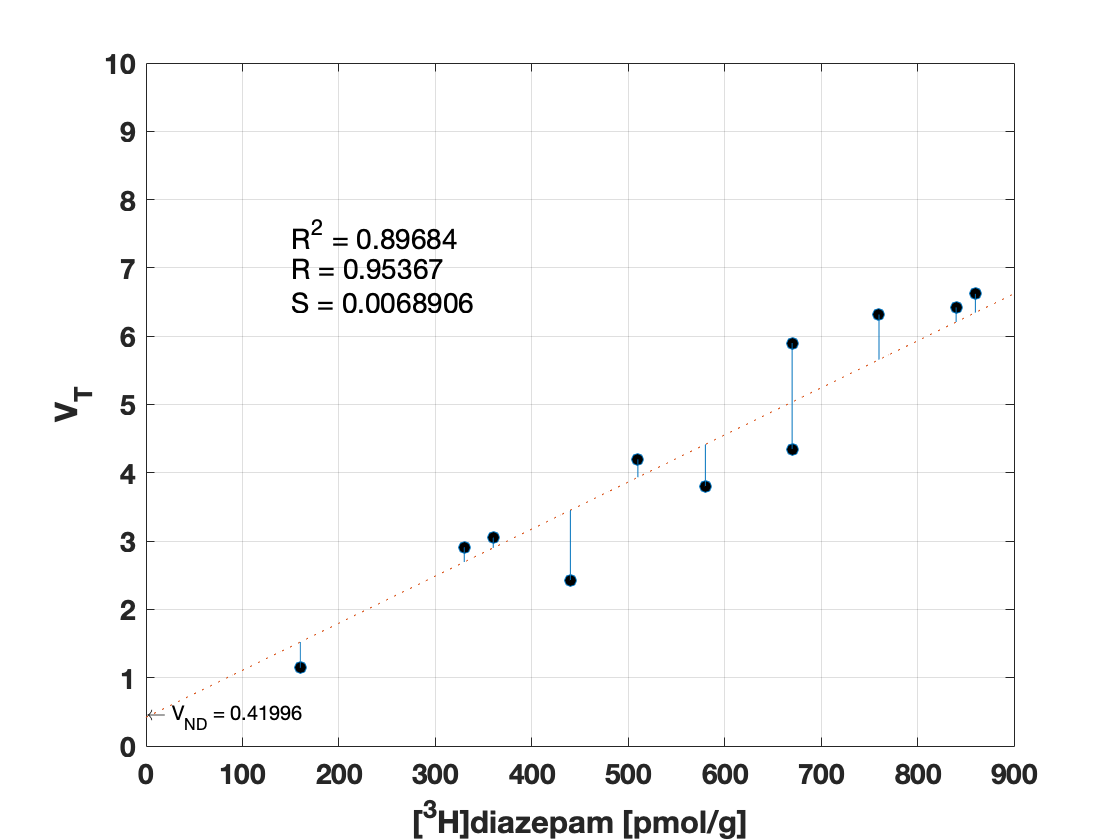

### Vt.png

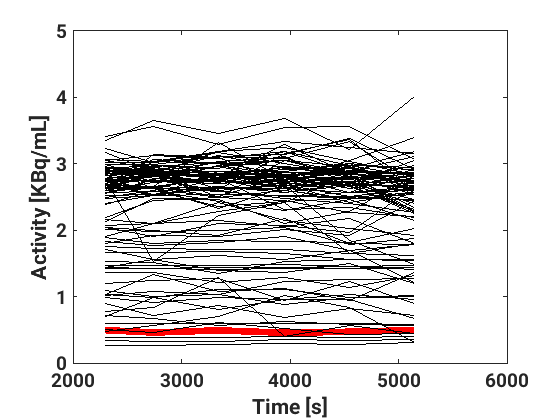

### Vt.png

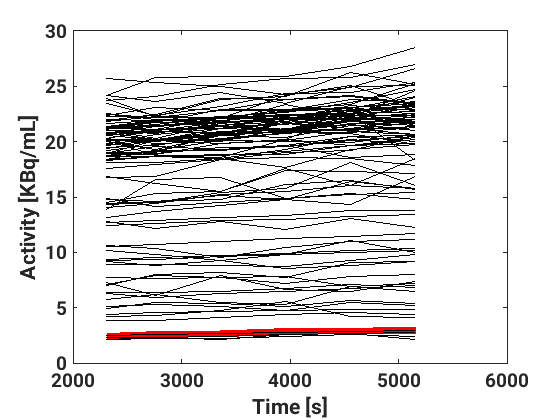

### Vt.png

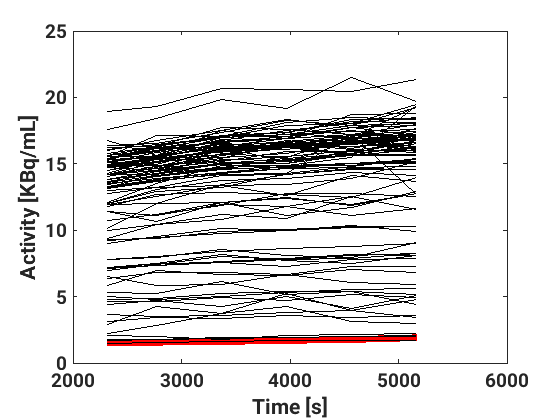

### Vt.png

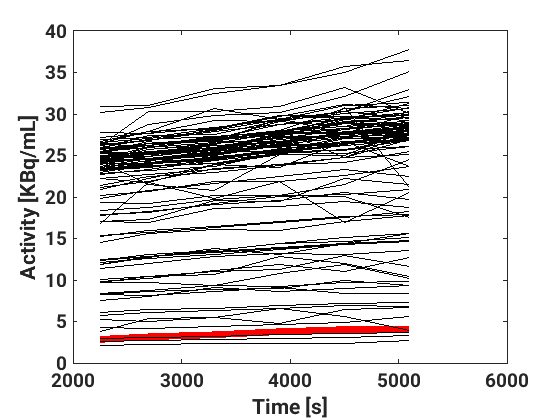

### Vt.png

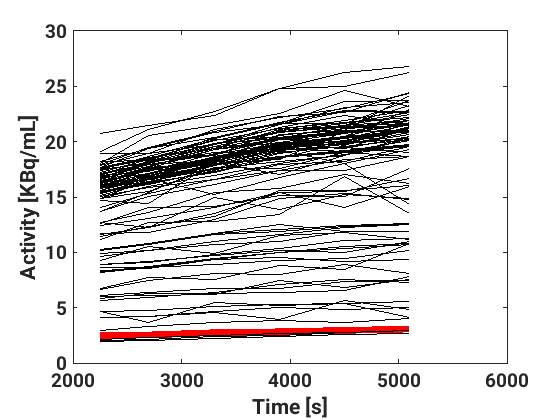

### Vt.png

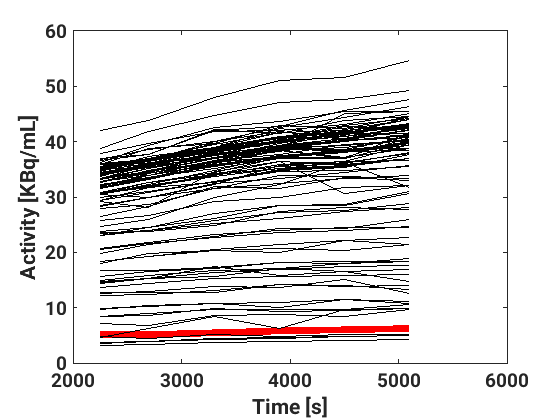

### Vt.png

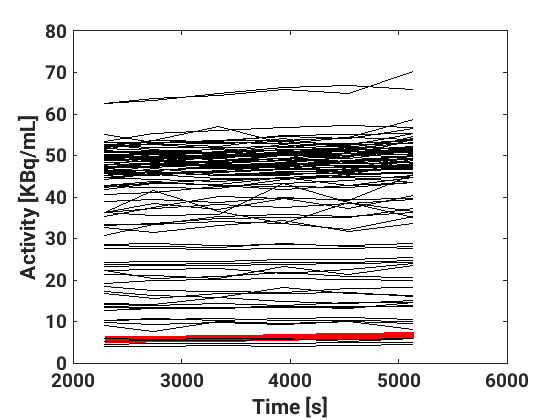

### Vt.png

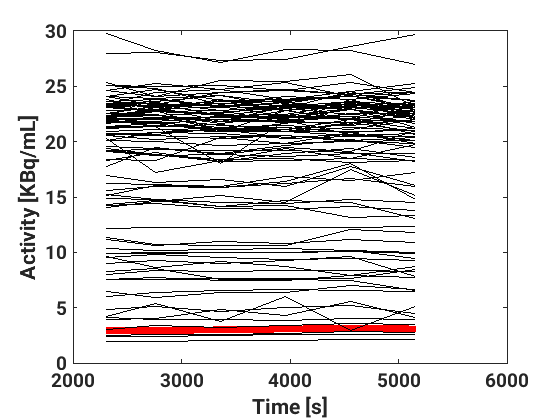

### Vt.png

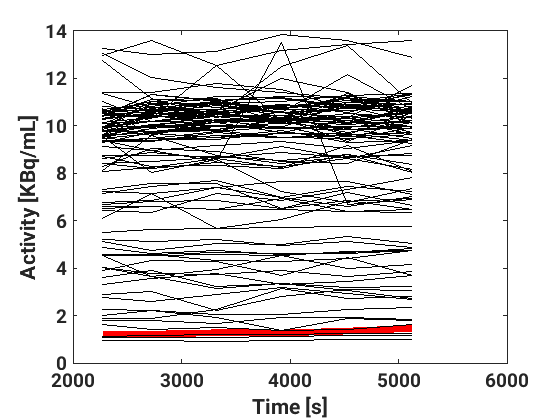

### Vt.png

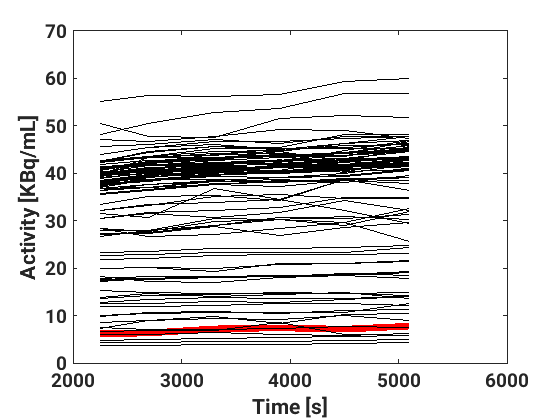

### Vt.png

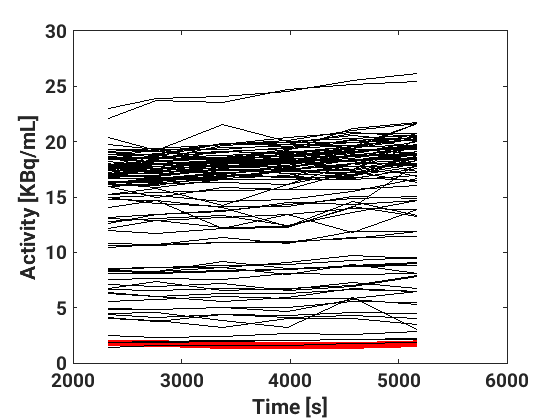

### Vt.png

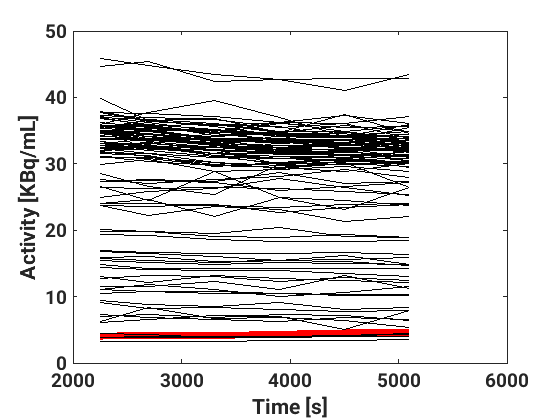

### Vt.png

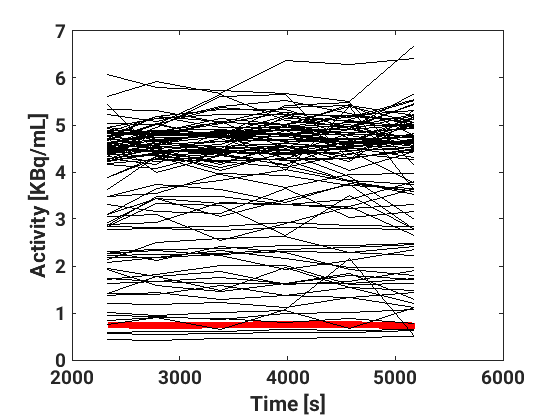

### Vt.png

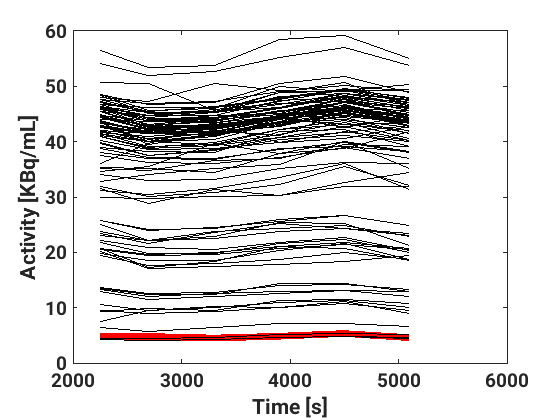

### Vt.png

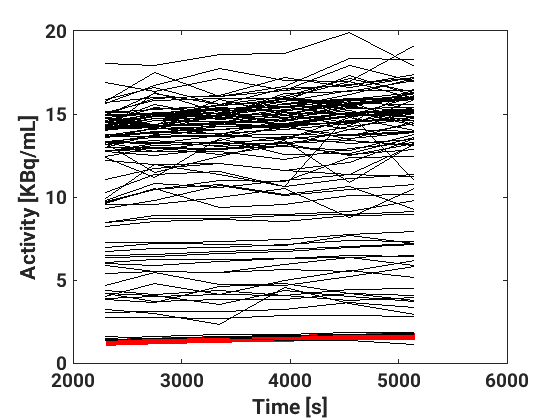

### Vt.png

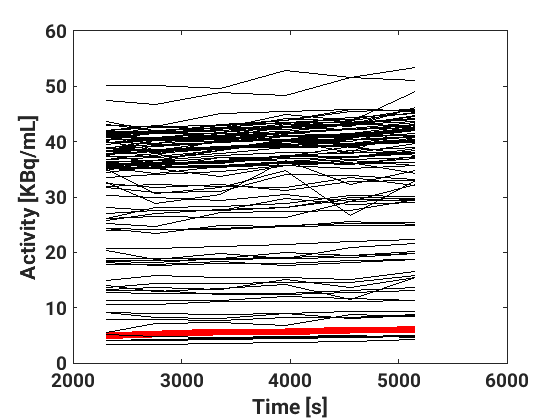

### Vt.png

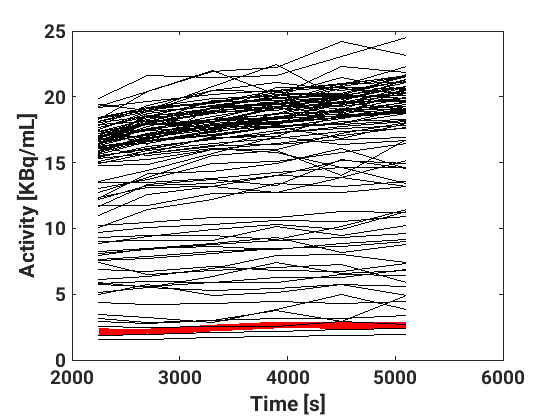

### Vt.png

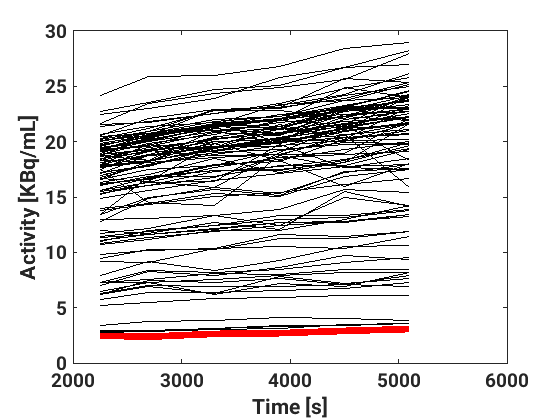

### Vt.png

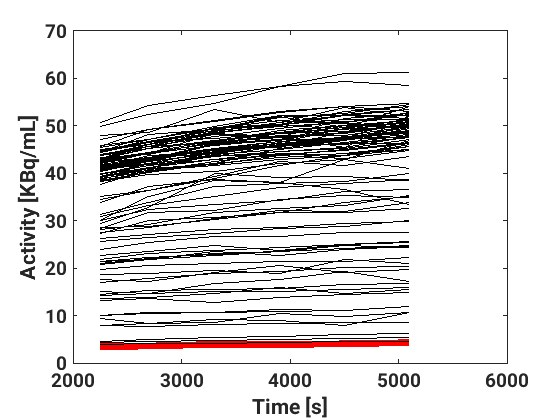

### Vt.png

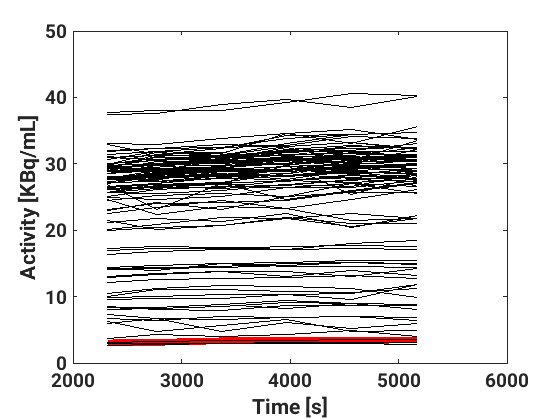

### Vt.png

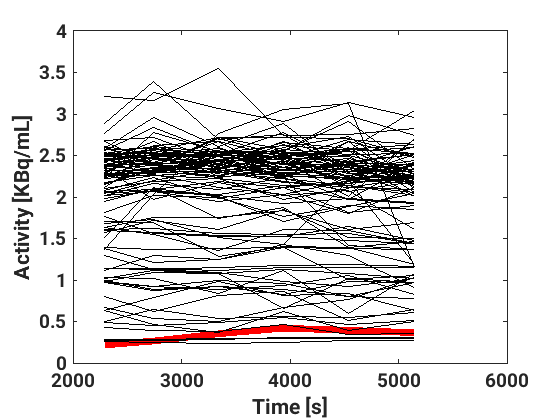
