## Supplementary material for "A High-Resolution In Vivo Atlas of the Human Brain’s Benzodiazepine Binding Site of GABA_A_ Receptors"

[2] University of Copenhagen, Faculty of Health and Medical Sciences, Copenhagen, Denmark

**Table S1:** PET scan protocols across subjects and scans

| | Bolus | Bolus-infusion | Individualized progarmmed infusion | Population-based programmed infusion |
|---|---|---|---|---|
| Subj 1 | | X (B/I=32m) | | |
| Subj 2 | | X (B/I=45m) | | |
| Subj 3 | | X (B/I=60m) | | |
| Subj 4 | | X (B/I=60m) | | |
| Subj 5 | X | | | |
| Subj 6 | X | | | |
| Subj 7 | X | | | |
| Subj 8 | | | | |
| Scan 1 | X | | | |
| Scan 2 | | | X | |
| Subj 9 | | | | |
| Scan 1 | | | X | |
| Scan 2 | | X (B/I=35m) | | |
| Subj 10 | | | | |
| Scan 1 | | X (B/I=35m) | | |
| Scan 2 | | | X | |
| Subj 11 | | | | |
| Scan 1 | X | | | |
| Scan 2 | | | X | |
| Subj 12 | | | | |
| Scan 1 | | | | X |
| Scan 2 | | X (B/I=55m) | | |
| Subj 13 | | | | |
| Scan 1 | | | | X |
| Scan 2 | | X (B/I=55m) | | |
| Subj 14 | | | | |
| Scan 1 | | | | X |
| Scan 2 | | X (B/I=55m) | | |
| Subj 15 | | | | |
| Scan 1 | | X (B/I=55m) | | |
| Scan 2 | | | | X |
| Scan 3 | | X (B/I=55m) | | |
| Subj 16 | | | | |
| Scan 1 | | X (B/I=55m) | | |
| Scan 2 | | | | X |
| **Total** | **5** | **12** | **4** | **5** |

**Table S2:** Regional distribution volumes ($V_T$) and non-displaceable distribution volume ($V_{ND}$) for each subject and scan.

| **Subj/Reg** | Fron | Cent | Occ | Temp | Cereb | Hippo | Amyg | Thal | Cau | Put | Pons | $V_{ND}$ |
|---|---|---|---|---|---|---|---|---|---|---|---|---|
| Subj 1 | 6.97 | 6.41 | 6.48 | 6.11 | 4.1 | 4.81 | 4.61 | 3.12 | 2.83 | 3.6 | 0.87 | 0.58 |
| Subj 2 | 8.06 | 7.39 | 7.3 | 7.0 | 4.37 | 5.14 | 5.41 | 3.56 | 2.85 | 4.08 | 1.09 | 0.66 |
| Subj 3 | 9.69 | 8.96 | 9.36 | 8.82 | 5.26 | 6.71 | 6.48 | 4.4 | 4.06 | 4.89 | 1.37 | 0.81 |
| Subj 4 | 6.3 | 5.89 | 5.83 | 5.55 | 3.4 | 4.21 | 4.01 | 2.6 | 2.1 | 2.95 | 0.87 | 0.26 |
| Subj 5 | 6.68 | 6.14 | 6.8 | 6.06 | 4.31 | 4.84 | 4.59 | 2.65 | 2.28 | 3.17 | 0.87 | 0.26 |
| Subj 6 | 7.29 | 7 | 6.56 | 6.24 | 3.86 | 5.14 | 4.89 | 2.95 | 2.79 | 3.36 | 0.91 | 0.34 |
| Subj 7 | 8.28 | 7.75 | 7.94 | 7.65 | 5.62 | 6.15 | 5.6 | 3.64 | 3.02 | 3.85 | 1.12 | 0.55 |
| Subj 8 | | | | | | | | | | | | |
| Scan 1 | 7.96 | 7.69 | 7.84 | 7.3 | 4.53 | 5.68 | 5.52 | 3.57 | 3.05 | 3.85 | 1.1 | 0.53 |
| Scan 2 | 8.09 | 7.84 | 7.7 | 7.21 | 4.43 | 5.52 | 5.43 | 3.28 | 2.96 | 3.69 | 0.98 | 0.17 |
| Subj 9 | | | | | | | | | | | | |
| Scan 1 | 8.03 | 8.14 | 7.49 | 6.38 | 4.59 | 6.44 | 6.47 | 4.03 | 3.62 | 4.62 | 1.43 | 1.31 |
| Scan 2 | 7.21 | 6.99 | 6.75 | 5.97 | 4.19 | 5.14 | 4.97 | 3.6 | 3.3 | 3.91 | 1.29 | 1.02 |
| Subj 10 | | | | | | | | | | | | |
| Scan 1 | 12.28 | 11.77 | 11.74 | 11.15 | 7.27 | 8.17 | 7.96 | 5.71 | 5.02 | 6.14 | 2.13 | 1.27 |
| Scan 2 | 7.71 | 7.16 | 8.02 | 7.17 | 4.77 | 5.52 | 5.36 | 3.76 | 3.18 | 3.93 | 1.33 | 0.8 |
| Subj 11 | | | | | | | | | | | | |
| Scan 1 | 7.55 | 7.41 | 7.58 | 6.81 | 4.31 | 5.23 | 4.93 | 3.02 | 2.67 | 3.21 | 1.0 | 0.12 |
| Scan 2 | 11.27 | 10.75 | 11.53 | 10.13 | 6.58 | 8.37 | 7.64 | 4.84 | 4.13 | 5.3 | 1.52 | 0.47 |
| Subj 12 | | | | | | | | | | | | |
| Scan 1 | 8.18 | 7.68 | 7.98 | 7.39 | 4.83 | 5.55 | 5.47 | 3.33 | 2.92 | 3.8 | 0.85 | 0.21 |
| Scan 2 | 10.47 | 9.95 | 10.39 | 9.58 | 6.1 | 7.32 | 6.89 | 4.35 | 4.0 | 5.01 | 1.0 | 0.29 |
| Subj 13 | | | | | | | | | | | | |
| Scan 1 | 9.77 | 9.12 | 9.66 | 8.79 | 5.9 | 7.1 | 6.78 | 4.56 | 4.04 | 4.78 | 1.24 | 0.75 |
| Scan 2 | 7.41 | 7.09 | 7.28 | 6.61 | 4.44 | 5.1 | 4.89 | 3.27 | 2.72 | 3.53 | 0.88 | 0.35 |
| Subj 14 | | | | | | | | | | | | |
| Scan 1 | 9.47 | 8.69 | 9.44 | 8.67 | 5.23 | 6.56 | 6.38 | 4.14 | 3.32 | 4.1 | 1.29 | 0.32 |
| Scan 2 | 8.5 | 7.91 | 8.31 | 7.63 | 4.64 | 5.75 | 5.46 | 3.52 | 2.82 | 3.55 | 0.96 | 0.04 |
| Subj 15 | | | | | | | | | | | | |
| Scan 1 | 7.78 | 7.2 | 7.73 | 7.18 | 4.88 | 5.08 | 5.11 | 3.39 | 3.02 | 3.83 | 1.25 | 0.58 |
| Scan 2 | 7.73 | 7.06 | 7.7 | 7.11 | 4.77 | 5.39 | 4.88 | 3.25 | 2.96 | 3.67 | 1.10 | 0.45 |
| Scan 3 | 8.3 | 7.6 | 8.34 | 7.55 | 5.11 | 5.69 | 5.2 | 3.57 | 3.18 | 3.93 | 1.14 | 0.42 |
| Subj 16 | | | | | | | | | | | | |
| Scan 1 | 6.06 | 5.76 | 6.02 | 5.47 | 3.48 | 4.18 | 3.99 | 2.82 | 2.39 | 3.21 | 1.03 | 0.6 |
| Scan 2 | 6.62 | 6.32 | 6.42 | 5.9 | 3.81 | 4.35 | 4.2 | 2.91 | 2.43 | 3.05 | 1.14 | 0.42 |
| **Average** | **8.22** | **7.76** | **8.01** | **7.36** | **4.8** | **5.73** | **5.5** | **3.61** | **3.14** | **3.96** | **1.15** | **0.52** |

**Table S3:** Linear mixed-effects model for $V_T$ ~ age+gender+age*gender+(1|administration)+(1|scan_number). The P-values were obtained using ANOVA with Kenward-Roger's method for estimating the degrees of freedom. All P-values are displayed *without* correction for mulitple comparisons using False-Discovery Rate (FDR). *With correction*, no P-values survive the 0.05 threshold, with the smallest P-values = 0.33 (range: 0.33-0.94).

| **Variable/Region** | Fron | Cent | Occ | Temp | Cereb | Hippo | Amyg | Thal | Cau | Put | Pons |
|---|---|---|---|---|---|---|---|---|---|---|---|
| Age | 0.88 | 0.66 | 0.93 | 0.94 | 0.82 | 0.61 | 0.55 | 0.55 | 0.84 | 0.84 | 0.49 |
| Gender | 0.06 | 0.09 | 0.04 | 0.03 | 0.06 | 0.12 | 0.11 | 0.24 | 0.45 | 0.27 | 0.62 |
| Age x Gender | 0.08 | 0.11 | 0.07 | 0.05 | 0.12 | 0.16 | 0.14 | 0.31 | 0.51 | 0.3 | 0.54 |

**Figure S1:** Flowchart of the processing of the MRI and PET data using the radioligand [$^{11}$C]Flumazenil, ranging from motion correction, matching of structural MRI using FreeSurfer (v. 6.0), kinetic modeling with arterial sampling, and finally establishing the association between postmortem human brain autoradiography from Braestrup et al. 1977 [4] and the regional total distribution volumes ($V_T$).

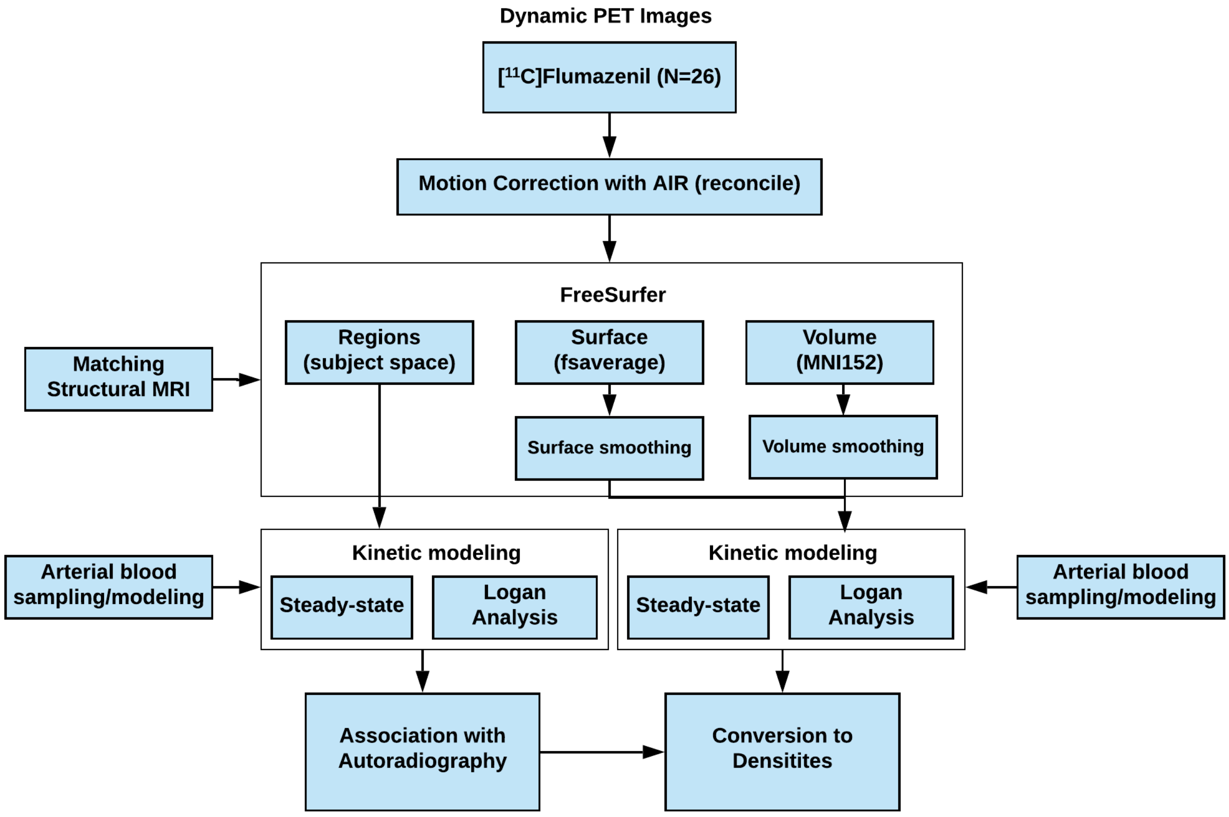

**Figure S2:** (left) Time activity curves for a single subject for several different brain regions using a bolus administration. (right) metabolite corrected blood input function. The distribution volume ($V_T$) for a given brain region was estimated using Logan with t* = 35 minutes. All fits of the Logan plot for a representative brain region (insula) can be found for all subjects in the supplementary folder (fit_blood_and_KinMod).

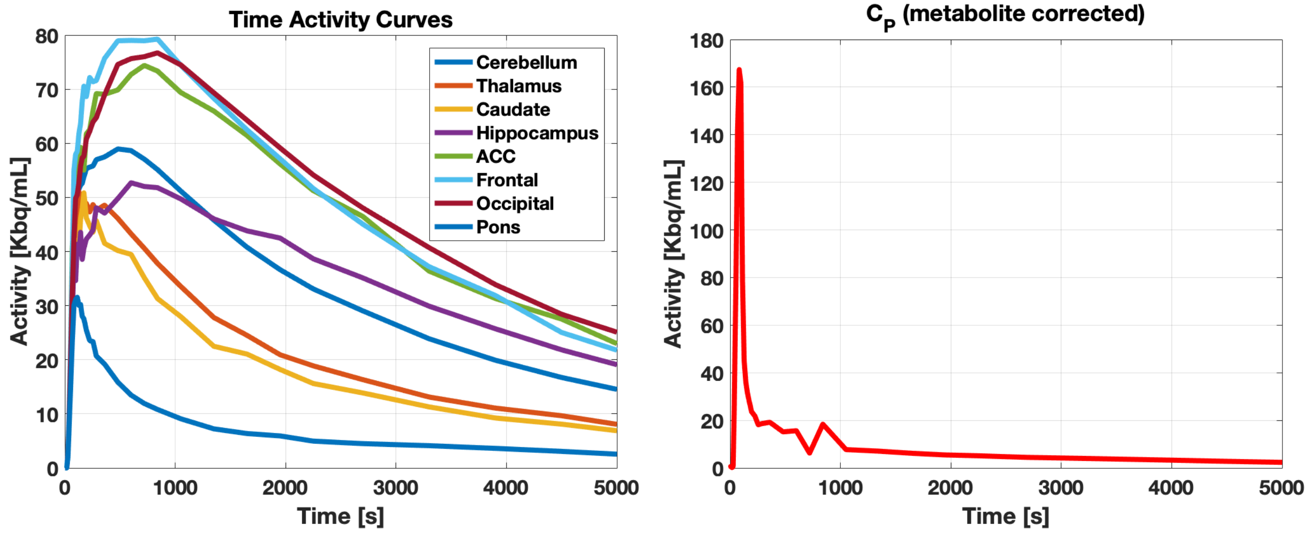


**Figure S3:** Time activity curves for a single subject for several different brain regions using a bolus-infusion administration. The red line is the metabolite corrected blood activity. This data can be found for all regions and subjects in the supplementary folder (fit_blood_and_KinMod).

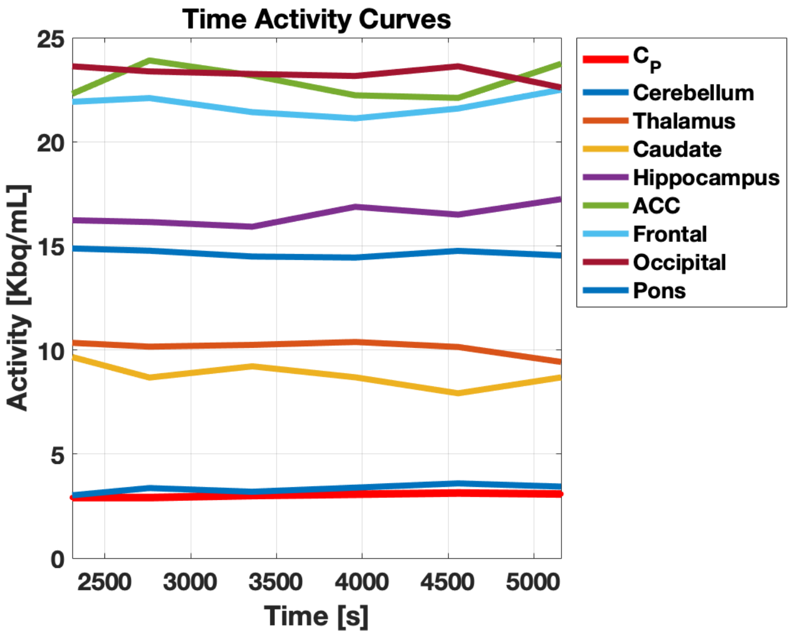

### Supplementary data:

**Bmax_vs_mRNA.xlsx:** Benzodiazepine receptor (BZR) density (pmol/ml) estimates for 48 brain regions (column 2). mRNA expression (log2 intensity) for 48 brain regions for each of the subunits of the $GABA_A$ receptor (column 3-21).

**Bmax.cvs.mni152.1mm.sm5.nii.gz:** Atlas of the benzodiazepine receptor (BZR) density (pmol/ml) in MNI152 space.

**Bmax.nopvc.fsaverage.lh.sm10.nii.gz:** Atlas of the benzodiazepine receptor (BZR) density (pmol/ml) of the left hemisphere in FSaverage space.

**Bmax.nopvc.fsaverage.rh.sm10.nii.gz:** Atlas of the benzodiazepine receptor (BZR) density (pmol/ml) of the right hemisphere in FSaverage space.

**Folder - correlation_mRNA_vs_Bmax:** Correlation plots of the benzodiazepine receptor (BZR) density (pmol/ml) estimates for 48 brain regions versus the corresponding mRNA expression for each of the 19 subunits of the $GABA_AR$.

**Folder - LinFit_autorad_vs_Vt:** Subject-specific linear fits of the autoradiography data from Braestrup et al. 1977 versus the distribution volumes obtained using PET.
